# A genetic screen to identify novel interactions of the phosphatidylinositol phosphate signaling pathway in Drosophila spermatogenesis

**DOI:** 10.64898/2026.09.25.754494

**Authors:** Sophie Rossini, Negar Nasirzadeh, Marynelle S. Icmat, Florence H-K. Chu, Mansuba Rana, Katarina Micovic, Kevinder Singh, Jingran Zhuo, Lisa Shao, Anamika Ohri, Kerui Huang, Maryam Lightwala, Murtaza S. Nagree, Tayyaba Jiwani, Kishan Bellamkonda, Preethy Prasad, Lilit Antonyan, Harrison Lands, Gianluca Sesso, Linda Liu, Nancy Fang Liu, Adrienne Philson, Dayag Sheykhkharimli, Priya Krithivasan, Andrew Papadopoli, Lacramioara Fabian, Julie A. Brill

**Author notes:** To whom correspondence should be addressed: Julie A. Brill, The Hospital for Sick Children PGCRL Building, 15^th^ Floor 686 Bay Street, Room 15.9716 Toronto, Ontario M5G 0A4 CANADA. Sophie Rossini, Negar Nasirzadeh, Marynelle S. Icmat, Florence H-K. Chu, Mansuba Rana, Katarina Micovic katarina.micovic@torontomu,.ca Kevinder Singh, Jingran Zhuo, Lisa Shao, Anamika Ohri, Kerui Huang, Maryam Lightwala, Murtaza S. Nagree, Tayyaba Jiwani, Kishan Bellamkonda, Preethy Prasad, Lilit Antonyan, Harrison Lands, Gianluca Sesso, Linda Liu, Nancy Fang Liu, Adrienne Philson,Dayag Sheykhkarimli, Priya Krithivasan, Andrew Papadopoli, Lacramioara Fabian, Julie A. Brill.

## Abstract

Phosphatidylinositol (PI) phosphates (PIPs) are membrane lipids with crucial roles in intracellular signaling, membrane trafficking and cytoskeletal organization. To perturb PIP function in *Drosophila melanogaster* sperm development, we previously expressed the Salmonella PIP phosphatase SigD (also known as SopB) under control of a spermatocyte-specific promoter. SigD expression results in reduced levels of plasma membrane PI 4,5-bisphosphate [PI(4,5)P2; henceforth PIP2] and male sterility. Transgenic lines expressing high levels of SigD (SigD-high) exhibit defects in spermatocyte cytokinesis, ciliary transition zone maturation and spermatid elongation, whereas lines expressing low levels of SigD (SigD-low) have defects in nuclear shaping and spermatid cell polarization. To further investigate the role of PIPs during Drosophila sperm development, we conducted a sensitized genetic screen to identify suppressors and enhancers of SigD-low phenotypes. From a screen of 760 chromosomal deficiencies spanning 83.6% of chromosomes 2 and 3, we identified 80 interacting regions that were further narrowed down to 40 individual suppressors. Among the strongest interactors were genes involved in microtubule dynamics, nucleocytoplasmic transport and membrane trafficking. The novel genetic interactions identified by our screen pave the way for further investigation into molecular mechanisms underlying the affected aspects of sperm development.

**Graphical Abstract:** 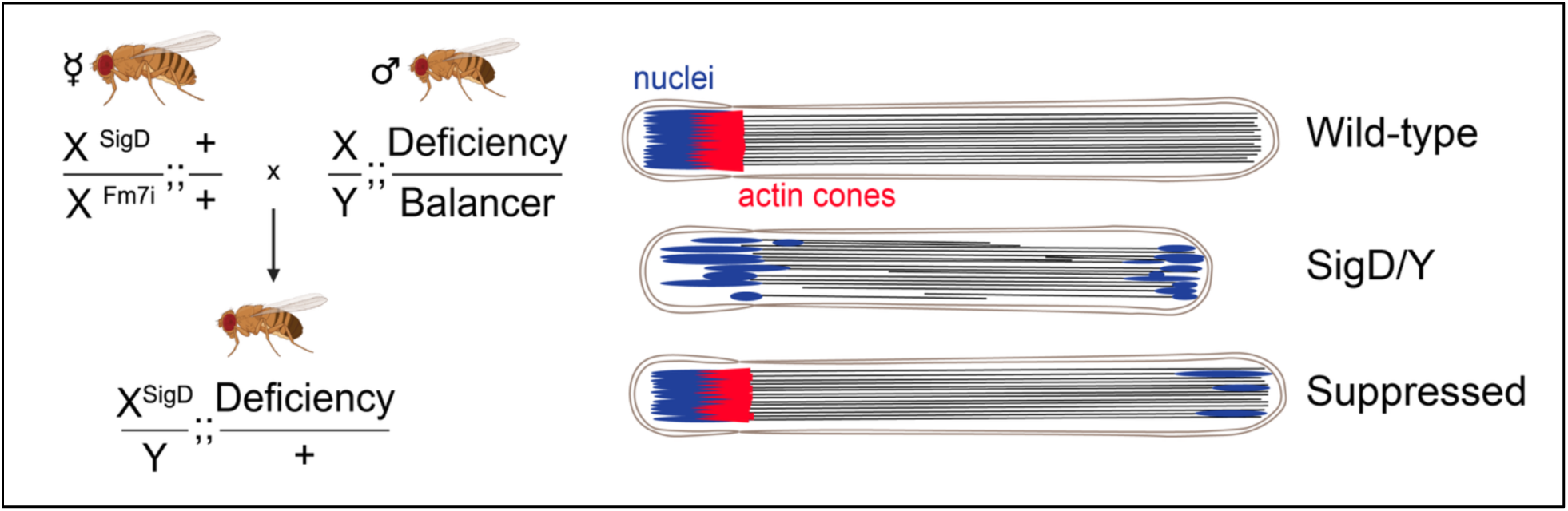

## Introduction

Spermatogenesis is the developmental process that produces sperm. Mammalian testes express >20,000 genes, including thousands of testis-enriched protein-coding genes (Soumillon et al. 2013; Djureinovic et al. 2014; Fagerberg et al. 2014; Uhlen et al. 2015). However, use of mammalian models to dissect genetic pathways contributing to sperm development is challenging. *Drosophila melanogaster* shares >2,000 testis-expressed genes with humans and provides a powerful system for performing screens to identify genes important for spermatogenesis (Wang et al. 2025).

In Drosophila, post-meiotic spermatids undergo extensive morphological changes to form mature sperm, a process called spermiogenesis (Fabian and Brill 2012). These changes include sperm head-tail coupling and elongation of sperm heads and tails (Rathke et al. 2007; Fabian and Brill 2012; Buglak et al. 2025). During elongation, all 64 spermatids polarize within a cyst: nuclei orient towards the base of the testis, and growing sperm tails point towards the testis tip (Fig. 1). At the end of elongation, an individualization complex (IC) consisting of 64 F-actin cones forms at the base of the spermatid nuclei and traverses the length of the sperm tails, removing unneeded cytoplasmic material (Steinhauer 2015). Upon reaching the sperm tail termini, the IC pinches off into a waste bag, and mature sperm are coiled and transferred to the seminal vesicle (Fabian and Brill 2012).

**Figure 1:**
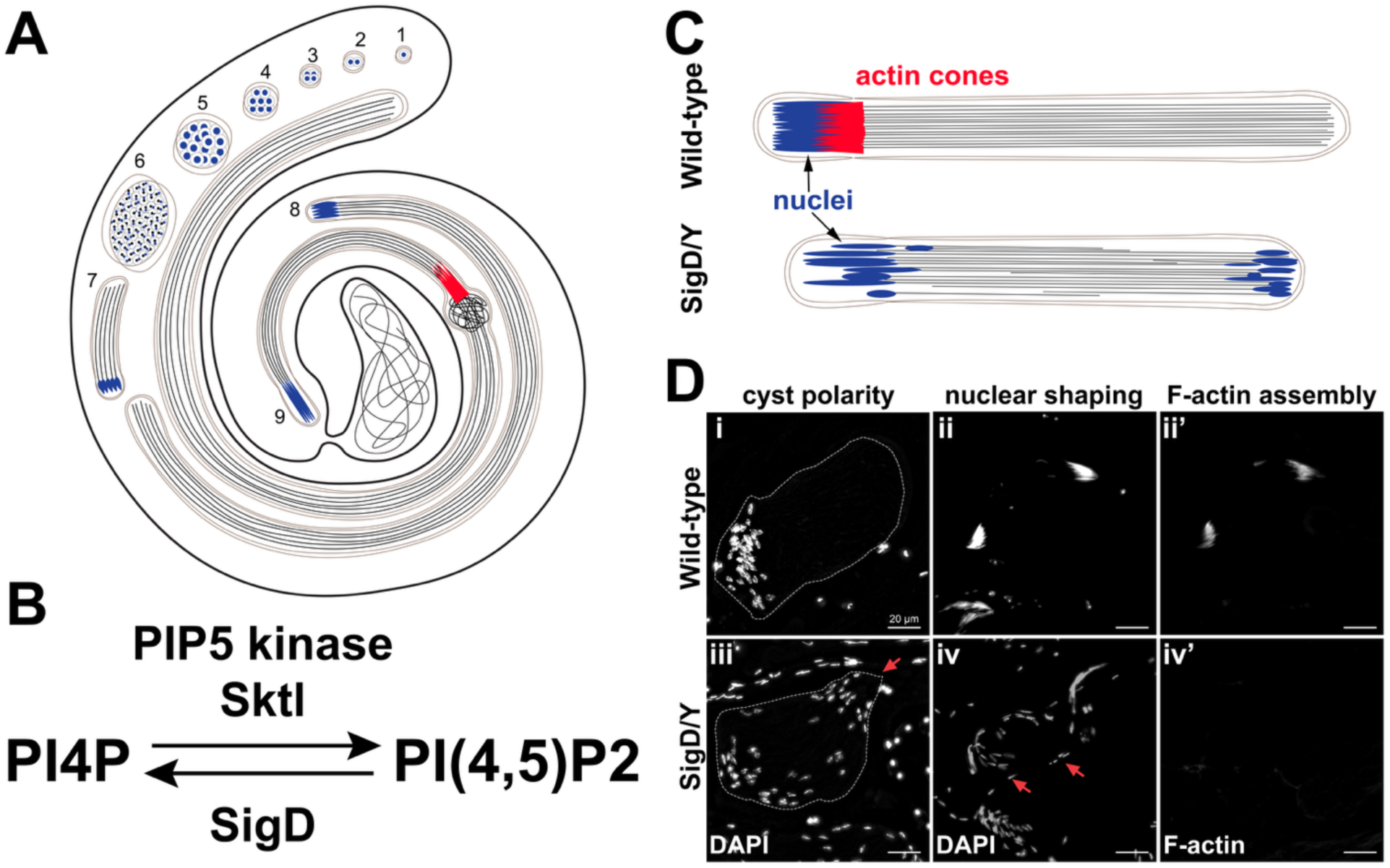
Drosophila spermatogenesis and SigD phenotypes. **(A)** Diagram of adult Drosophila testis showing stages of spermatogenesis. Spermatogenesis begins at the tip of the testis, where a germline stem cell (GSC; Fig. 1A.1) divides asymmetrically to yield a new GSC and a gonialblast that is committed to differentiation. The gonialblast undergoes four mitotic amplification divisions with incomplete cytokinesis (Fig. 1A.2-5) to give rise to a cyst of 16 interconnected spermatogonia, which undergo pre-meiotic S-phase to become primary spermatocytes. Spermatocytes grow 25-fold in volume and express ∼8,000 genes before undergoing meiosis I and II (Fig. 1A.6) to produce a cyst of 64 interconnected round spermatids that undergo spermiogenesis to form mature sperm (Fig. 1A.7-9). Blue: nuclei, red: actin cones. **(B)** PIP5-kinases, such as Sktl in Drosophila, phosphorylate PI4P to generate PI(4,5)P2 (PIP2). Exogenous expression of the Salmonella effector protein SigD results in dephosphorylation of PIP2 to PI4P. **(C-D)** Testes of wild-type males possess unipolar cysts and tightly clustered needle-shaped nuclei and actin cone complexes. Loss of PIP2 in SigD-low male germ cells results in bipolar cysts (red arrow, left), scattered and misshapen nuclei (red arrows, middle), and complete loss of individualization complexes containing actin cones.

Male germ cell development relies on signaling by phosphatidylinositol phosphates (PIPs), membrane phospholipids that are phosphorylated by lipid kinases at the -3, -4, or -5 positions of the inositol head group (Fabian and Brill 2012; Brill et al. 2016). The PIP 5-kinases that synthesize PIP2 from PI 4-phosphate (PI4P) are essential for sperm development in flies and mice (Wei et al. 2008; Fabian et al. 2010; Hasegawa et al. 2012; Gupta et al. 2018). We previously generated transgenic Drosophila lines expressing SigD (also called SopB), a Salmonella effector protein that dephosphorylates PIP2 to PI4P or interconverts PIP2 to PI(3,4)P2 (Steele-Mortimer et al. 2000; Terebiznik et al. 2002; Walpole et al. 2022), under control of the spermatocyte-specific *β_2_-tubulin* (*β_2_t*) promoter. SigD expression reduces plasma membrane PIP2 levels and produces infertile male flies (Wei et al. 2008; Fabian et al. 2010). SigD was expressed at either high or low levels, depending on whether the transgene was inserted on an autosome (SigD-high) or the X-chromosome (SigD-low). SigD-high lines exhibit severe spermatogenesis defects, including multinucleate spermatids and delayed spermatid cyst elongation (Wong et al. 2005; Wei et al. 2008). SigD-low lines show milder phenotypes, including bipolar cysts (nuclei at both ends), misshapen nuclei, and failure to form actin cones during spermiogenesis (Fabian et al. 2010) (Fig. 1). Notably, both high and low levels of SigD abrogate sperm head-tail coupling (Wei et al. 2008; Fabian et al. 2010). The milder SigD-low phenotypes provide a sensitized background to screen for large deletions (deficiencies, Dfs) and genes whose heterozygous loss modifies phenotypes associated with reduced PIP2 levels. Here, we describe the results of this screen, which identified 40 genetic suppressors of phenotypes associated with SigD-low. Our data reveal novel PIP2 pathway interactors, including potential regulators or targets of PIP2 signaling during sperm development.

## Materials and methods

### Fly stocks and husbandry

Flies were raised on standard cornmeal molasses agar at 25°C (Ashburner 1990). Transgenic flies expressing low levels of SigD under control of the spermatocyte-specific *β_2_t* promoter were previously described (Wong et al. 2005; Wei et al. 2008; Fabian et al. 2010). Because the SigD-low transgene is inserted on the X-chromosome, deficiency kit stocks spanning chromosomes 2 and 3 (Table S1) were obtained for screening from the Bloomington Drosophila Stock Center. *w^1118^* was used as the wild-type control.

### Primary screen

Virgin *w^1118^,* P{*w^+^, β_2_t-SigD*}*/FM7i, w^-^*balanced females were crossed to male flies from individual balanced Bloomington deficiency stocks (male flies of genotype *Df*/+, where “+” indicates a second- or third-chromosome balancer). Testes from 1-2 day-old P(*w^+^*) male progeny were dissected and stained for DNA and F-actin using 1:1000 DAPI (VWR 89139-054) and 1:400 rhodamine phalloidin (Invitrogen R415), as described (Hime et al. 1996). Testes were examined on an upright Zeiss Axioplan-2 epifluorescence microscope using a 40x objective lens, and images were acquired using Axiovision software and an Axiocam black and white camera (Carl Zeiss).

### Secondary screen

Once the interacting intervals were defined (Table S2), alleles of individual genes within each interval were tested for their ability to suppress or enhance phenotypes caused by SigD-low. Males carrying alleles of interest (as homozygotes or heterozygous over balancer chromosomes) were tested as described above for the Df lines. All interactors were tested independently over a period of several years by at least four lab members, with the strongest 40 interactors being selected for further validation. Stocks used for individual alleles are found in Table S3.

### Molecular Cloning

To generate a construct expressing *mRuby3*-tagged *skittles* (*sktl*, which encodes the Drosophila PIP 5-kinase Sktl that synthesizes PIP2) in Drosophila testes, the NEBuilder website (https://nebuilderv1.neb.com/) was used to generate primers to amplify *mRuby3* and linker::*sktl* (Invitrogen). PCR fragments were amplified using Phusion^TM^ High-Fidelity DNA Polymerase (NEB M0530L) with the following primers: *mRuby3* 5’-TTTTTCGGTGTGAAAAAGGGATGGTGTCTAAGGGCGAAG-3’ and 5’-GCGGCCGCCCCTTGTACAGCTCGTCCATG-3’, linker::*sktl* 5’-GCTGTACAAGGGGCGGCCGCAGATGTCC-3’ and 5’-GGTTCCTTCACAAAGATCCTTCTAGACCTCAACGCGGCAAATCG-3’. The *β_2_t* vector tv3(Wong et al. 2005) was digested using *Xba*I and *EcoR*I (New England Biolab, NEB) according to manufacturer’s instructions. PCR fragments and digested *β_2_t* vector were run on a 1% agarose gel, and DNA was extracted using GenepHlow^TM^ Gel/PCR Kit (Geneaid DFH300). NEBuilder^®^ HiFi DNA Assembly Cloning Kit (NEB E5520) was used for Gibson assemblies according to the manufacturer’s guidelines. The assembled plasmid was transformed into NEB DH5-alpha competent *E. coli* (NEB C2987) using a standard heat shock transformation protocol and carbenicillin selection. Plasmid DNA was extracted using Monarch® Spin Plasmid Miniprep Kit (NEB T1110L). The *β_2_t-mRuby3::sktl* plasmid was confirmed by Sanger sequencing (TCAG, The Hospital for Sick Children) and integrated into the attP40 site on the second chromosome via PhiC31 integrase-mediated transgenesis by injection into *y^1^ w^67c23^; P{CaryP}attP40* embryos (BestGene Inc. Chino Hills, CA, USA). To carry out genetic experiments described below, the *y^1^ w^67c23^* X-chromosome was replaced with *w^1118^*. As above, standard Gibson cloning was used to add an *mNeonGreen* (mNG) tag to the C-terminus of the PIP2-binding domain *PLCδ-PH* (De Vos et al. 2003). *PLCδ-PH* was amplified using the following primers: 5’-CCTAGAGGATCCCCGGGTACCATGGACTCGGGCCGGGAC-3’ and 5’-CCATGGTGGCGACCGGTGGATCCTTCAGGAAG-3’. *mNG* was amplified using the following primers: 5’-TCCACCGGTCGCCACCATGGTGAGCAAG-3’ and 5’-TCTATTTGGCTTTAGTCGCTCTAGATTACTTGTACAGCTCGTCCATG -3’. *PLCδ-PH* and *mNG* were then cloned into a P-element vector containing the *α_1_-tubulin* promoter cut with *Kpn*I and *Xba*I. The *α_1_-tub-PLCδ-PH::mNG* plasmid was confirmed by Sanger sequencing (TCAG, The Hospital for Sick Children). Transgenic flies were generated by injection of *w^1118^* embryos, using as a helper the Δ2,3 transposase-expressing plasmid, pTurbo (Laski et al. 1986; Robertson et al. 1988). Transgenic lines were mapped and balanced using *w^1118^*; *Sco/CyO* to generate *w^1118^*; P{*w^+^, α_1_-tub-PLCδ-PH::mNG*}/*CyO*.

### Testis Immunostaining

Testis immunostaining was performed as described (Hime et al. 1996). Briefly, testes of newly eclosed (1-2 day old) male flies were dissected in testis isolation buffer (TIB) (Casal et al. 1990), cut with tungsten needles, and squashed with coverslips on Polysine^TM^ Microscope Adhesion Slides (ThermoFisher Scientific P4981-001). Samples were placed in chilled 95% ethanol for a minimum of 10 min, then fixed using 4% paraformaldehyde (Electron Microscopy Science 15710) for 7 min, then briefly washed with PBT (PBS and 0.1% Triton X-100), followed by two 15 min washes in PBT-DOC (PBS, 0.1% Triton X-100, and 0.3% sodium deoxycholate). After two 5 min washes with PBT, samples to be immunostained were blocked for 30 min in PBTB (PBT and 5% bovine serum albumen) and incubated overnight at 4° C with 1:1000 anti-acetylated α-tubulin (Sigma-Aldrich T7451). The next day, samples were washed with PBT three times for 5 min, then incubated at room temperature for 1 hr with 1:1000 goat anti-mouse Alexa Fluor^TM^ 488 (Invitrogen A11001). Following fixation and permeabilization (with or without immunostaining), samples were washed for 15 min in PBT, incubated with 1:400 Alexa Fluor^TM^ 594 Phalloidin (Invitrogen A12381) or rhodamine phalloidin (Invitrogen R415) for 30 min, washed twice in PBT for 15 min each, and mounted using ProLong^TM^ Glass Antifade Mountant with NucBlue^TM^ (Invitrogen P36985).

### Live Imaging

To image live squashed preparations of Drosophila spermatocytes expressing the PIP2 marker PLCδ-PH-mNG, testes of newly eclosed (1-2 day old) male flies were dissected in TIB with 1:1000 Hoechst 33342 solution (20 mM) (ThermoFisher Scientific 62249) to stain DNA. Samples were cut with tungsten needles and squashed with coverslips on Polysine^TM^ Microscope Adhesion Slides (ThermoFisher Scientific P4981-001) before viewing on a Leica DMi8 epifluorescence microscope.

### Microscopy

Phase-contrast and fluorescence images of live or fixed samples were observed using a 40x objective on an inverted Leica DMi8 microscope equipped with an LED3 light source, Leica K5 camera and THUNDER Imaging System. Images were acquired and adjusted for brightness and contrast using LAS-X software (Version 3.7.4.23463, Leica Microsystems Ltd., Wetzlar, Germany) and ImageJ (Fiji Version 2.1.0/1.53c). All figures were created using Adobe Illustrator (Version 25.4.1 (2021), Adobe Inc., San Jose, CA, USA). Unless indicated, control and experimental samples were imaged under identical conditions, and images were adjusted only for contrast and brightness using identical manipulations.

## Results

### Deficiency screening identifies interacting genomic intervals and candidate genes

To identify genes that interact with the PIP pathway, a deficiency (Df) modifier screen was performed using SigD-low as a sensitized background. Examination of SigD-low testes stained for F-actin and DNA by fluorescence microscopy confirmed defects in sperm head-tail coupling, spermatid polarization, nuclear elongation, and actin cone formation (Fabian et al. 2010) (Fig. 1C-D). We hypothesized that heterozygous deletion of regions and genes that interact with PIPs would suppress or enhance SigD-low phenotypes.

To screen large genomic regions, we used the Drosophila deficiency (Df) kit (Cook et al. 2012), which contains a set of large deletions (Dfs) that remove ∼84% of chromosomes 2 and 3. Because the SigD-low transgene is on the X, and males have only one X chromosome, we performed an F_1_ genetic modifier screen using Dfs that span regions of chromosomes 2 and 3. To generate “modified” males, female flies carrying the SigD-low transgene marked by red eyes (*w^+^*) were crossed to male flies harboring individual Dfs. F_1_ males carrying the SigD-low transgene (*w^+^*) and one copy of the Df were collected and screened for suppression or enhancement of spermatid polarization, nuclear elongation, or actin cone defects. We screened a total of 760 Bloomington Dfs (Fig. 2, Table S1), of which 75 resulted in suppression and 5 resulted in enhancement of the SigD-low phenotypes (Table S2).

**Figure 2.**
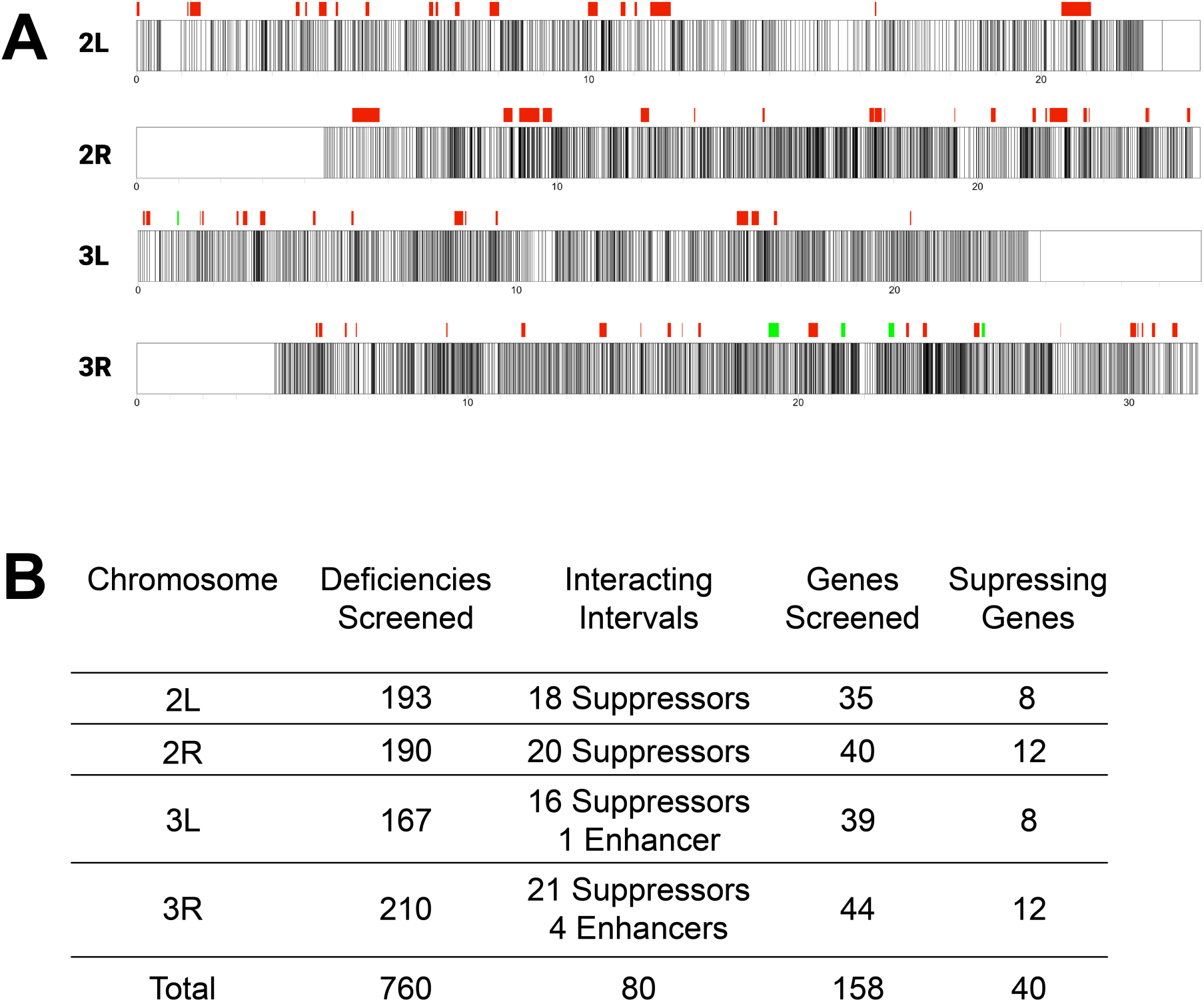
Summary of deficiency screen to find genetic interactors of the PIP pathway. **(A)** Map of chromosomes 2 and 3 where relative positions of interacting suppressors and enhancers in red and green respectively are shown. **(B)** A screen of 760 deficiencies from the Bloomington Deficiency kit, covering 83.6% of chromosomes 2 and 3, identified 75 suppressing (red) and 5 enhancing (green) genomic intervals. From these interacting intervals, ostensible null or hypomorphic alleles of 134 genes were screened. Of these 134 genes, alleles of 40 genes resulted in genetic suppression, and no alleles resulted in genetic enhancement, of phenotypes caused by expression of SigD.

To further refine regions of interest, we compared the effects of smaller deficiencies that were contained within, or partially overlapped with, the larger Dfs. We reasoned that if a smaller Df produced the same level of phenotypic suppression or enhancement as the larger Df, it likely contained the gene(s) responsible for the interaction identified in the initial screen (Fig. 3A). Conversely, smaller Dfs that did not interact genetically with SigD-low were presumed to lack the relevant gene(s). It is also possible that simultaneous loss of multiple genes (rather than a single candidate) within a larger Df is required for the interaction, an aspect that our screening strategy would not adequately capture. After narrowing the regions of interest, we tested mutant alleles of individual genes within the identified regions for suppression or enhancement of SigD-low phenotypes. Although we found no single gene responsible for enhancement of SigD-low phenotypes, we identified 99 genes that individually suppressed SigD-low (Table S3), including 40 that showed relatively strong suppression (Figure 3B, Table S4). Of these, three genes—*CG2955*, *Nup98-96, and TSG101*—exhibited particularly strong, reproducible suppression phenotypes.

**Figure 3.**
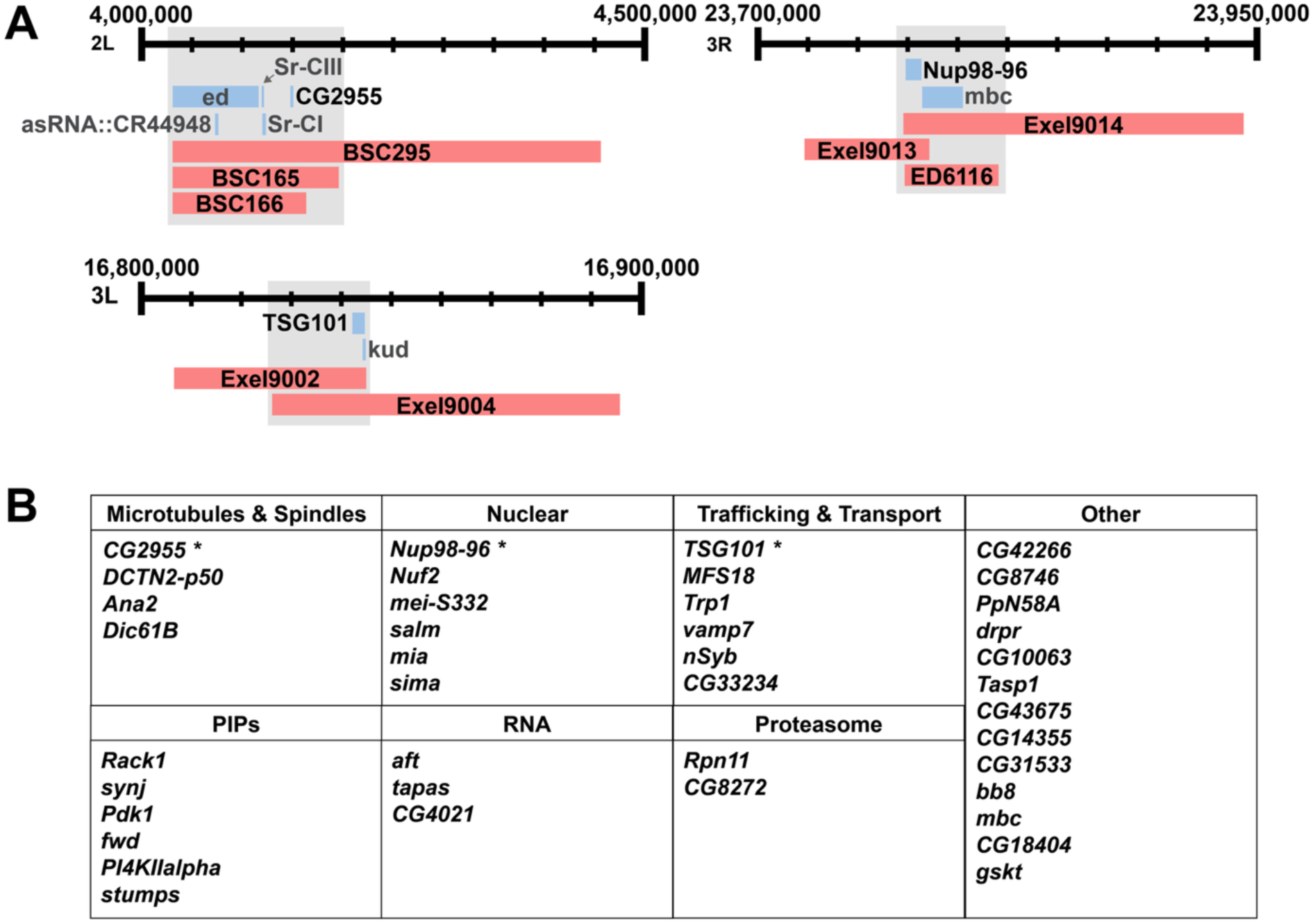
Genomic intervals of the top three strongest SigD suppressors. **(A)** Candidate genes (light blue) were identified by narrowing down genes found within overlapping suppressing deficiencies (red). Hypomorphic or null alleles of each candidate gene were tested for suppression of cyst polarity, nuclear shaping, or actin cone formation. The strongest three suppressors (black text)—*CG2955*, *Nup98-96,* and *TSG101*—suppressed phenotypes caused by reduced levels of PIP2. **(B)** The 40 suppressors were categorized into seven groups based on their established functions. The three strongest suppressors (*) are associated with pathways related to microtubule dynamics, nucleocytoplasmic transport or membrane trafficking.

### Heterozygous loss of CG2955, Nup98-96, or TSG101 suppresses SigD-low phenotypes

To characterize suppression of SigD-low, we examined sperm tails, (acetylated α-tubulin), as well as sperm heads (DNA) and actin cones (F-actin) in testes from wild-type control flies, SigD-low flies, and SigD-low flies heterozygous for mutant alleles of each of *CG2955*, *Nup98-96*, or *TSG101* (Fig. 4, 5). As a positive control, we also assessed testes of flies expressing an additional copy of *mRuby3::sktl* to rescue the loss of PIP2 caused by SigD expression. In wild-type control testes, we observed the expected morphology of spermatid cysts: early elongated cysts were polarized, with nuclei at one end and sperm tails growing in the opposite direction; at later stages, the nuclei of mature sperm were elongated and needle-shaped; onset of individualization was marked by formation of the IC at the base of elongated nuclei (Fig. 4A-A’, 5A-A’’). In contrast, SigD-low spermatids formed bipolar cysts with nuclei at both ends of the cyst and, in some cases, with nuclei scattered throughout the cyst. The nuclei were not elongated or only partially elongated, and the F-actin cones did not form (Fig. 4B-B’, 5B-B’’). Co-expression of *mRuby3::sktl* partially rescued all these defects: cysts were more polarized; more nuclei localized at one end of the cyst; some nuclei were able to achieve needle-shape; and the IC formed at the base of the elongated nuclei (Fig. 4C-C’, 5C-C’’). Although infertility was not suppressed, the extent of suppression by co-expression of *mRuby3::sktl* supports a role for PIP2 in spermiogenesis.

**Figure 4.**
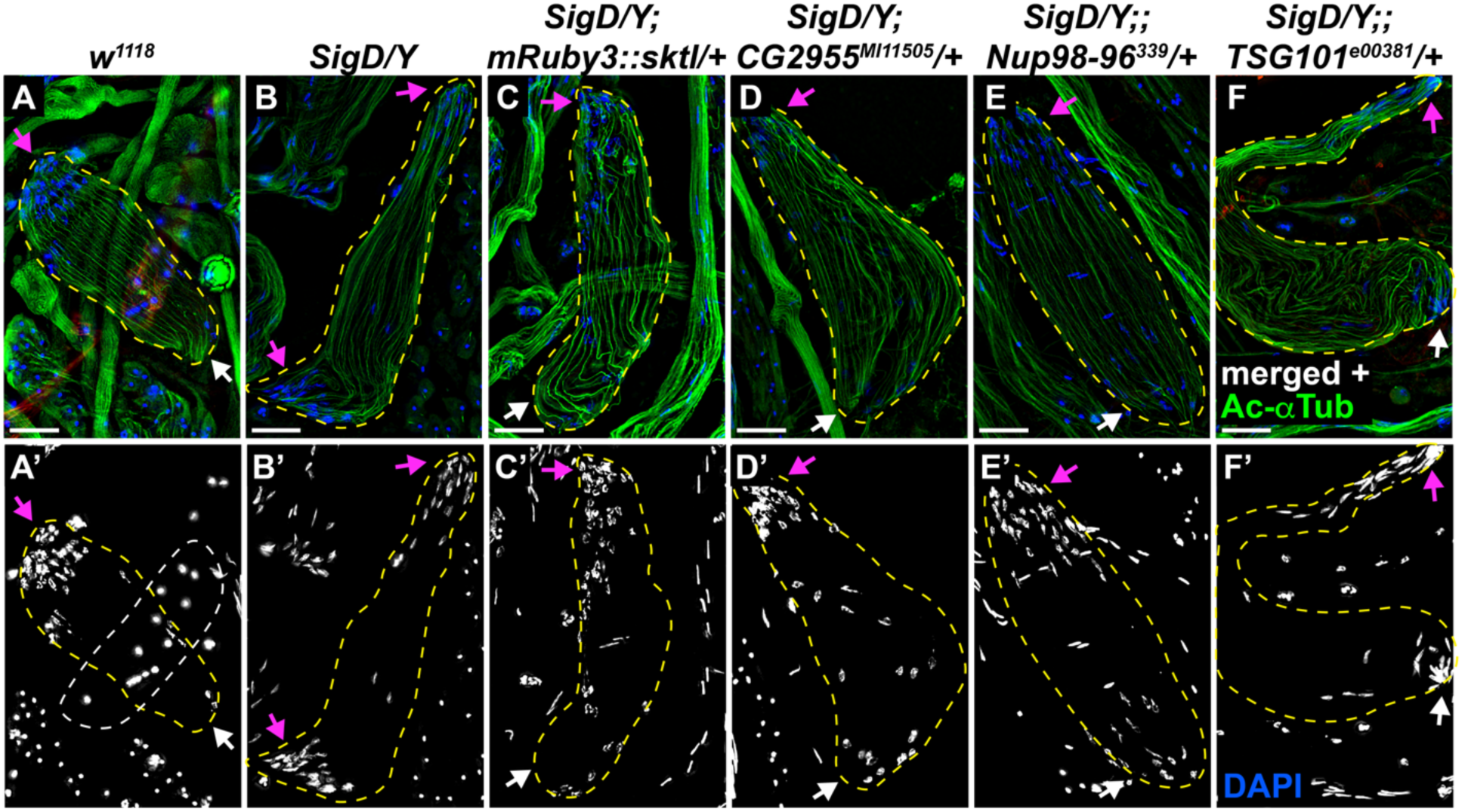
Partial suppression of SigD spermatid polarization defects. Epifluorescence micrographs showing localization of acetylated α-tubulin (green) and DNA (blue) in early elongating spermatid cysts. Individual cysts are traced with dashed yellow lines. In control (*w^1118^*) **(A),** nuclei were found at only one end of the cyst (compare magenta vs. white arrows), whereas in the presence of SigD **(B)**, nuclei were found at both ends of the cyst. The spermatid polarization defect was partially suppressed by co-expression of *mRuby3::sktl* **(C)**, as evidenced by the higher concentration of nuclei at one end of the cyst, as compared to SigD expression alone (compare magenta vs. white arrows in C’ vs. B’). Heterozygous loss of *CG2955* **(D)**, *Nup98-96* **(E)**, or *TSG101* **(F)** partially suppressed the spermatid polarization defects in SigD. *CG2955* was a stronger suppressor of the cyst polarity defect than *Nup98-96* or *TSG101*. White dashed line indicates nuclei from an underlying cyst in A’. Data shown are representative of more than 3 separate experiments. Scale bar: 20 µm.

**Figure 5.**
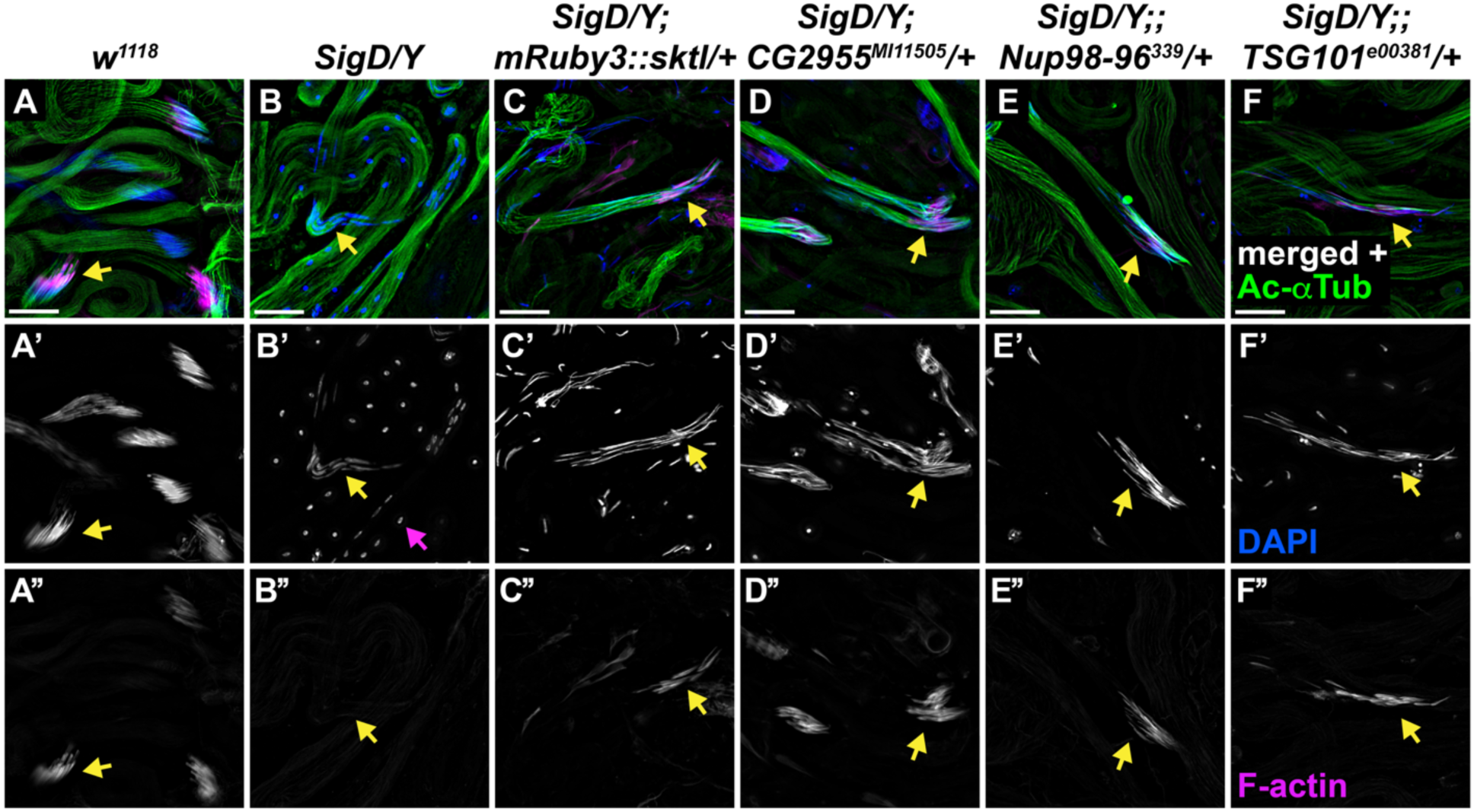
Suppression of nuclear elongation and F-actin cone formation defects in SigD spermatids. Epifluorescence micrographs showing localization of sperm tails (anti-acetylated α-tubulin, green), F-actin (rhodamine phalloidin, magenta) and nuclei (DAPI, blue) in elongated spermatid cysts. In control (*w^1118^)* **(A)**, all 64 spermatid nuclei clustered at one end, and the elongated nuclei were needle-shaped (yellow arrows). F-actin cones appeared at the base of the elongated nuclei, indicating onset of individualization. In SigD **(B),** a majority of the nuclei were scattered throughout the cyst (magenta arrow), with a few cysts progressing to mid-elongation (yellow arrow). SigD nuclei never became needle-shaped, and actin cones never formed at the base of the spermatid nuclei. Co-expression of *mRuby3::sktl* **(C)** partially suppressed the nuclear defects in SigD; the nuclei became needle-shaped (yellow arrow in C’) and the individualization complex formed at the base of the elongated nuclei (yellow arrow in C”). Heterozygous loss of *CG2955* **(D)**, *Nup98-96* **(E)**, or *TSG101* **(F)** suppressed the nuclear elongation defects, as evidenced by the presence of needle-shaped nuclei and actin cones. Data shown are representative of more than 3 separate experiments. Scale bar: 20 µm.

SigD-low flies heterozygous for *CG2955, Nup98-96,* and *TSG101* mutant alleles demonstrated partial suppression of SigD phenotypes, albeit not to the extent of testes expressing an additional copy of *mRuby3::sktl*. Notably, *CG2955* was the strongest suppressor of cyst polarity, showing the most dramatic rescue of spermatid polarization among the three suppressors (Fig. 4D-F’, 5D-F’’)*. CG2955* encodes an uncharacterized microtubule (MT) end binding protein paralog with enriched expression in testes compared to other tissues (Vedelek et al. 2018). Thus, our study is the first to provide evidence that *CG2955* plays a role in proper progression of spermiogenesis.

*Nup98-96* and *TSG101* have previously been implicated in Drosophila spermatogenesis. Loss of both copies of *Nup98-96* results in loss of germline stem cells and disruption of spermatogonial mitotic divisions during aging (Parrott et al. 2011). In contrast, a previous study knocking down *TSG101* in somatic cyst cells, but not germline cells, results in overproliferation of early germline cells and spermatogonial tumorigenesis (Tang et al. 2017). Our data suggest additional roles for *Nup98-96* and *TSG101* during later stages of sperm development.

### Heterozygous loss of CG2955, Nup98-96 or TSG101 does not rescue PIP2 levels in SigD-low

To determine if suppression of SigD phenotypes by heterozygous loss of *CG2955, Nup98-96*, or *TSG101* is due to restoration of PIP2 levels, we assessed PIP2 levels in spermatocytes, the cells where SigD is first expressed, using the fluorescent PIP2 marker PLCδ-PH (Varnai and Balla 1998) fused to mNeonGreen. As expected, PIP2 localized at the plasma membrane (PM) of wild-type spermatocytes (Fig. 6A-A’). In cells expressing SigD-low, PIP2 was nearly undetectable at the PM, whereas co-expression of *mRuby3::sktl* with SigD-low restored PM PIP2 (Fig. 6B-C’). Loss of one copy of *CG2955*, *Nup98-96*, or *TSG101* in a SigD background did not restore PIP2 levels, as no observable signal was detected at the spermatocyte PM (Fig. 6D-F’). Thus, heterozygous loss of these genes did not promote PIP2 synthesis.

**Figure 6.**
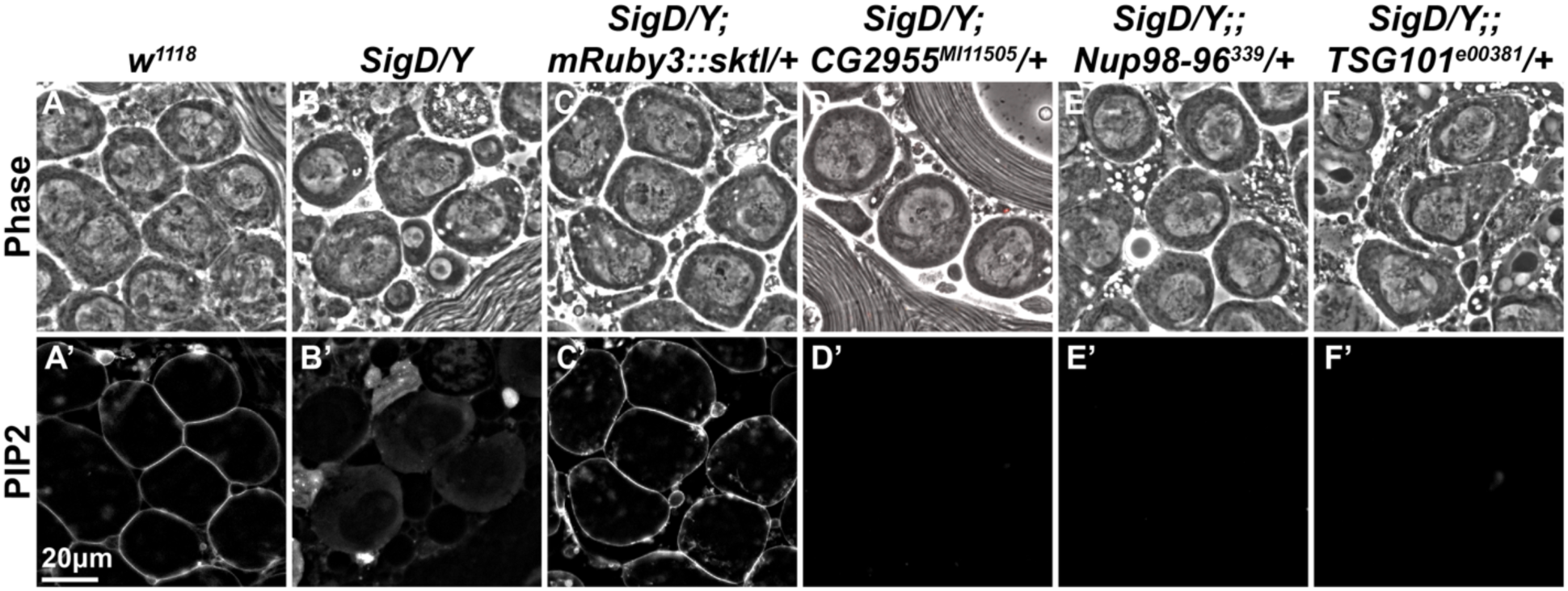
PIP2 levels are not restored in SigD-low spermatocytes heterozygous for the suppressors. Phase contrast (upper panel) and epifluorescence micrographs (lower panel) of spermatocytes expressing PLCδ-PH-mNG marking PIP2. In control (*w^1118^)* **(A)**, PIP2 localized robustly at the plasma membrane (PM) in spermatocytes. Expression of SigD-low **(B)** resulted in dramatically reduced levels of PM PIP2 and increased cytoplasmic levels of PLCδ-PH-mNG. Co-expression of *mRuby3::sktl* **(C)** with SigD restored PM PIP2, whereas heterozygous loss of *CG2955* **(D)**, *Nup98-96* **(E),** or *TSG101* **(F)** in the SigD background did not restore PM PIP2 levels. Scale bar: 20 µm.

## Discussion

In a sensitized genetic screen, we identified 40 genes that suppressed phenotypes caused by PIP2 depletion in developing Drosophila male germ cells. Among these suppressors were genes involved in microtubule dynamics, nucleocytoplasmic transport, membrane trafficking, transcription, ubiquitination, and mRNA binding. The strongest suppressors (*CG2955*, *Nup98-96*, *TSG101*) are linked to diverse cellular processes—microtubule dynamics, nucleocytoplasmic transport and membrane trafficking, suggesting these processes intersect with PIP signaling to drive successful spermiogenesis in Drosophila.

*CG2955* encodes a testis-specific MT plus-end binding (EB) protein. *CG2955* null mutants are viable and fertile, yet loss of one copy of *CG2955* suppresses effects of PIP2 depletion, suggesting some of the defects observed in SigD might be due to effects on MT polymerization or stability at the nuclear envelope. Loss of one copy of *CG2955* may correct this defect, allowing for successful nuclear shaping. A paralog of *CG2955*, *Mst27D*, bundles MTs during nuclear elongation (Li et al. 2023). It is currently unclear how loss of one copy of *CG2955* improves spermiogenesis in PIP2-depleted cells. However, loss of *Mst27D* prevents nuclear elongation, and loss of one copy of *CG2955* promotes nuclear elongation, suggesting *CG2955* and *Mst27D* may have opposite effects on MT function in this process.

*Nup98-96* is essential for transit-amplifying divisions of spermatogonia (Parrott et al. 2011), yet its role in spermiogenesis is unknown. The allele used in this study specifically affects Nup96, which is part of Nup107-160, a nine protein subcomplex of the nuclear pore complex. Notably, GCP3 a core member of γ-TuRC specifically interacts with Nup96 to promote MT nucleation at kinetochores (Mishra et al. 2010). This interaction may also contribute to MT organization during spermiogenesis, where Nup96 may interact with γ-TuRC to generate MTs required during sperm nuclear elongation, in addition to its canonical role in nucleocytoplasmic transport.

Tumour susceptibility gene 101 (*TSG101*) encodes a core component of the ESCRT-I complex, which mediates endocytic recycling and endolysosomal trafficking of PM proteins and is involved in cell polarization (Moberg et al. 2005; Sevrioukov et al. 2005; Giebel and Wodarz 2006; Lobert and Stenmark 2011). Spermatid elongation involves a ∼150-fold increase in length, and individualization requires extensive membrane remodelling leading to PM surrounding each individual sperm. These morphogenic events likely place substantial demands on membrane trafficking pathways. Although its role in spermiogenesis is unclear, heterozygous loss of *TSG101* partially suppressed defects in F-actin assembly and spermatid polarization caused by PIP2 depletion, implicating ESCRT-mediated trafficking in these processes.

Drosophila sperm development is a complex process that depends on the functions of >1000 proteins (Dorus et al. 2006; Wasbrough et al. 2010; Rettie and Dorus 2012), and many of the pathways contributing to spermatogenesis remain unknown. Through a sensitized genetic screen, we identified multiple suppressors of defects in PIP2 signaling, revealing an intimate relationship between the encoded proteins and PIP2 signaling. Because heterozygous loss of these proteins did not directly increase PIP2 levels, we surmise that these genes identify pathways that modulate spermiogenesis in a manner that depends on PIP2. Overall, by uncovering spermatogenesis mechanisms in Drosophila, we may begin to elucidate genetic pathways fundamental to sperm development and male fertility in humans.

## Acknowledgements

We are grateful to research volunteers Mallika Makkar, Jing-Yi Yuan, Sonya Snyder and Spencer Brown for assistance with the screen; Paul Paroutis and Kimberly Lau (SickKids Imaging Facility) for fluorescence microscopy training; the Bloomington, Kyoto and Harvard Drosophila Stock Centers, and colleagues Thomas Schwarz (Boston Children’s Hospital, Massachusetts, USA), Andre Bedard (McMaster University, Ontario, Canada), Christian Lehner (Universität Zürich, Switzerland), Frank Schnorrer (Institut du Biologie du Développement de Marseille, France), Stephan Sigrist (Freie Universität Berlin, Germany), Helen White-Cooper (Cardiff University, Wales), Ernst Hafen (ETH Zürich, Switzerland), Marc Freeman (Vollum Institute, Oregon Health Sciences University, Portland, USA), Shelagh Campbell (University of Alberta, Canada), Mark Van Doren (Johns Hopkins University, Baltimore, USA), Barbara Wakimoto (University of Washington, Seattle, USA), Paul Schedl (Princeton University, USA), Carmen Robinett (Janelia Research Campus, Ashburn, USA), William Sullivan (University of California Santa Cruz, USA), Eduardo Moreno (Institute of Cell Biology, Universität Bern, Switzerland), James Kennison (Section on Gene Regulation, National Institute of Child Health and Human Development, NIH, Bethesda, USA), Armen Manoukian (Princess Margaret Hospital, Toronto, Ontario, Canada) for Drosophila stocks; and Flybase version FB2026_02 for crucial information about Drosophila genes identified in our screen.

---

**Supplementary Table 1a.**
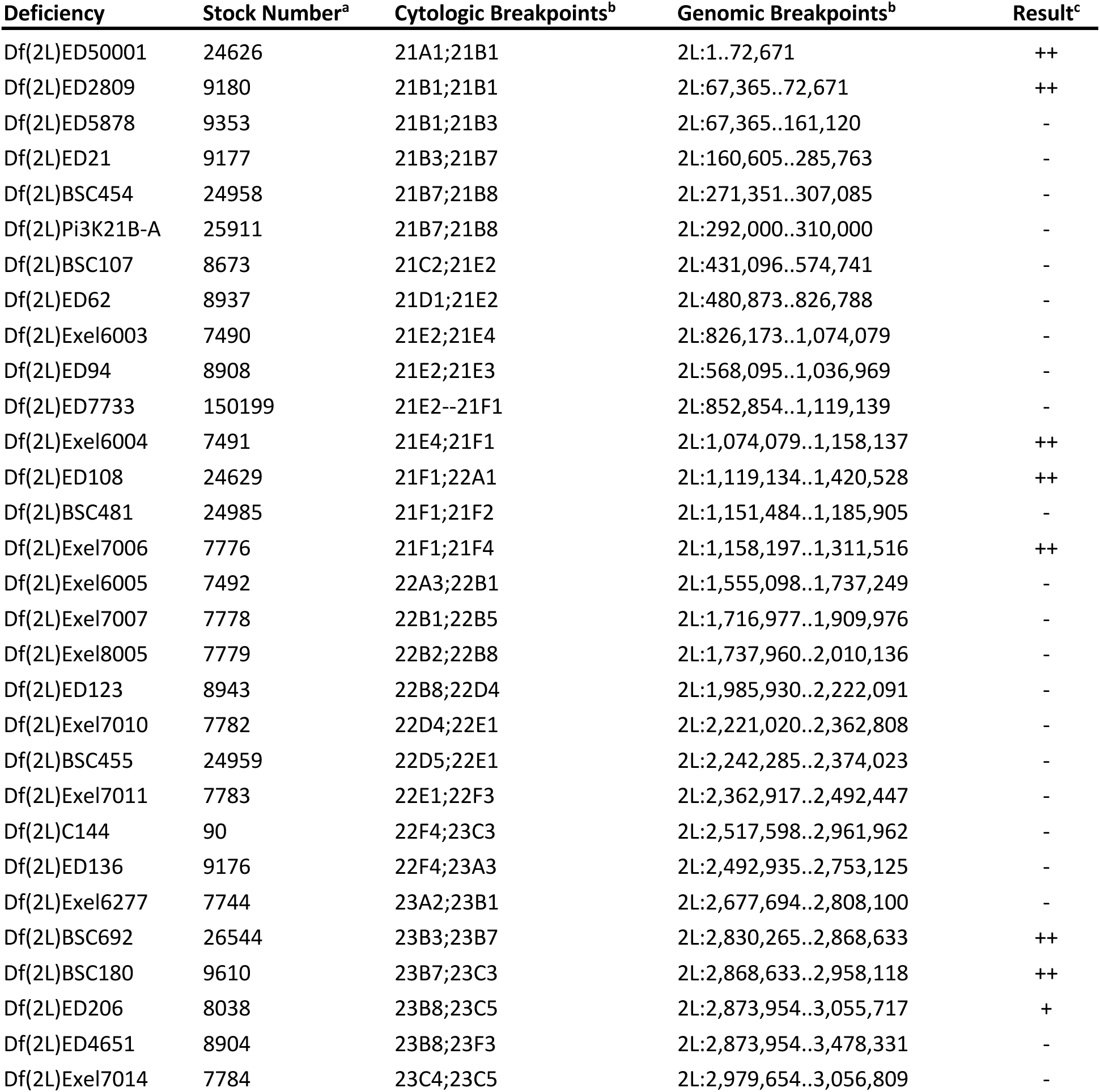
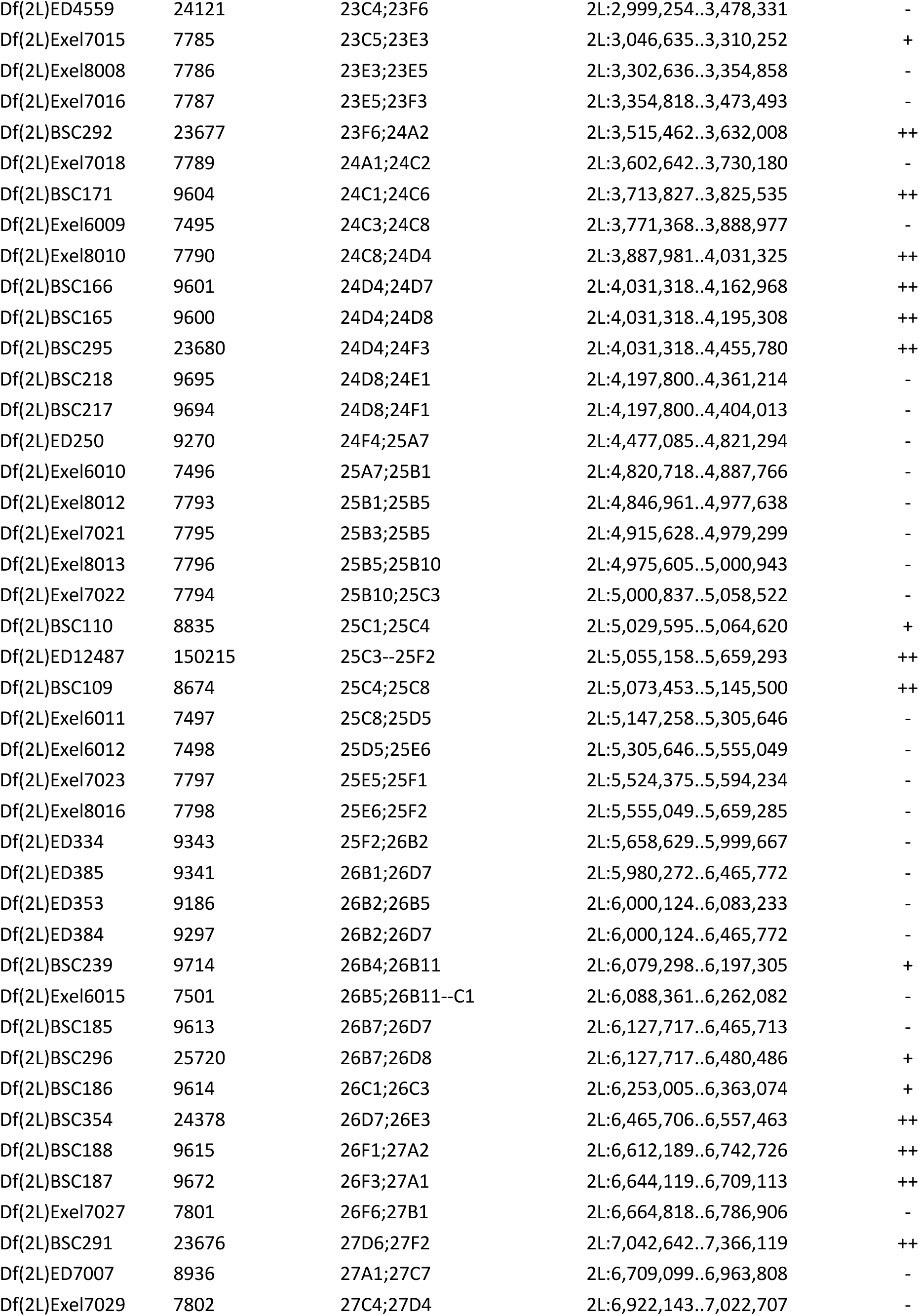
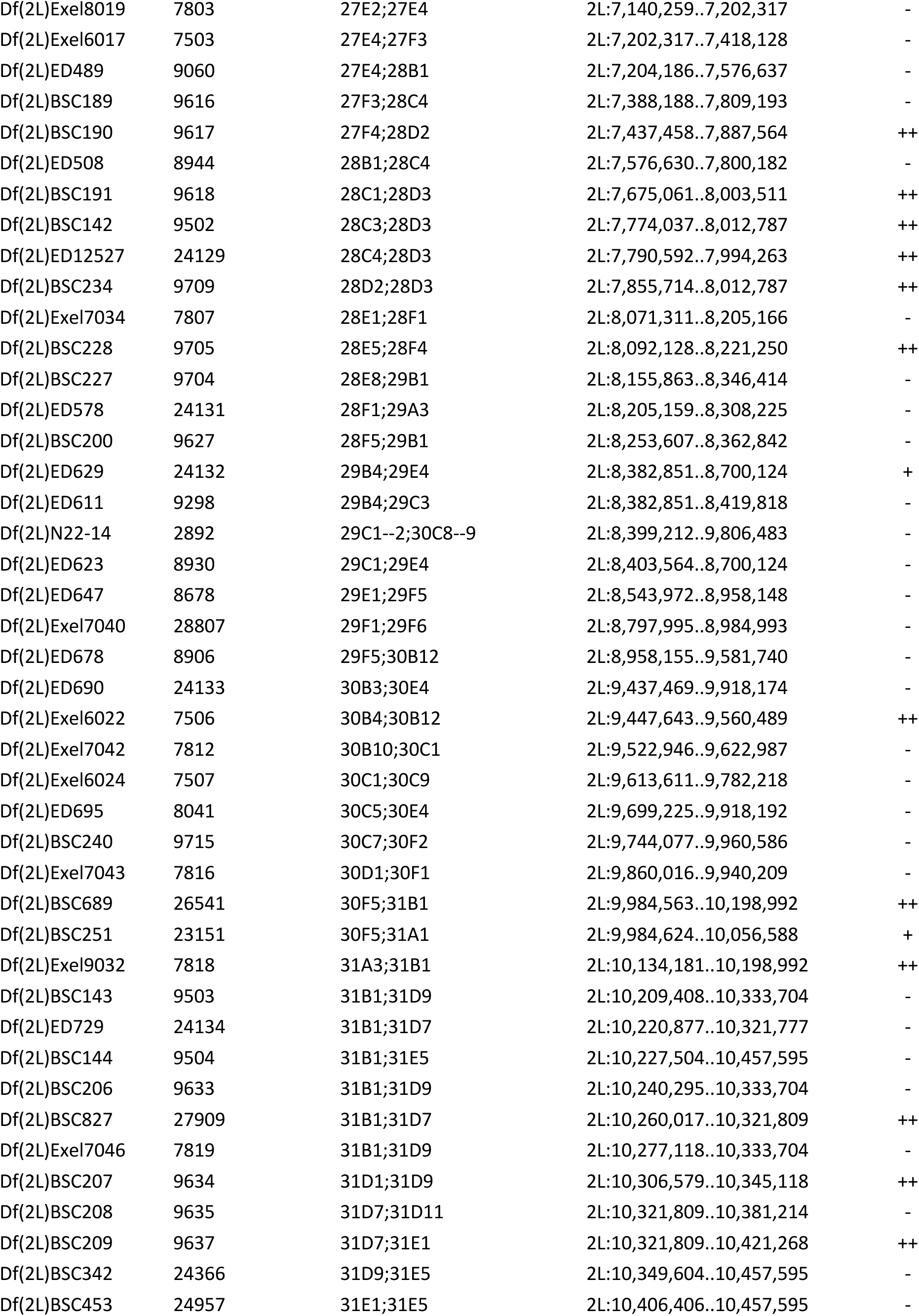
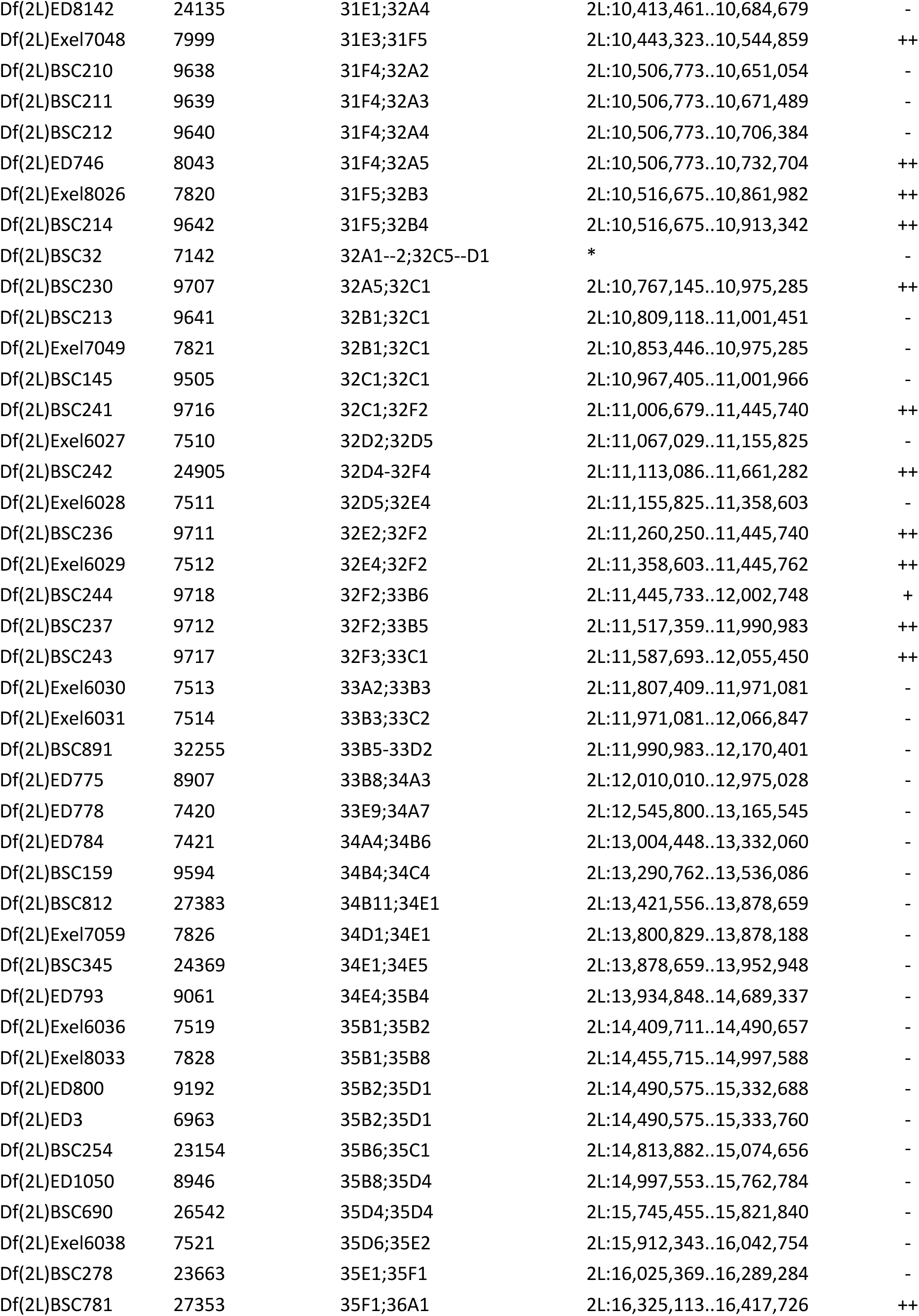
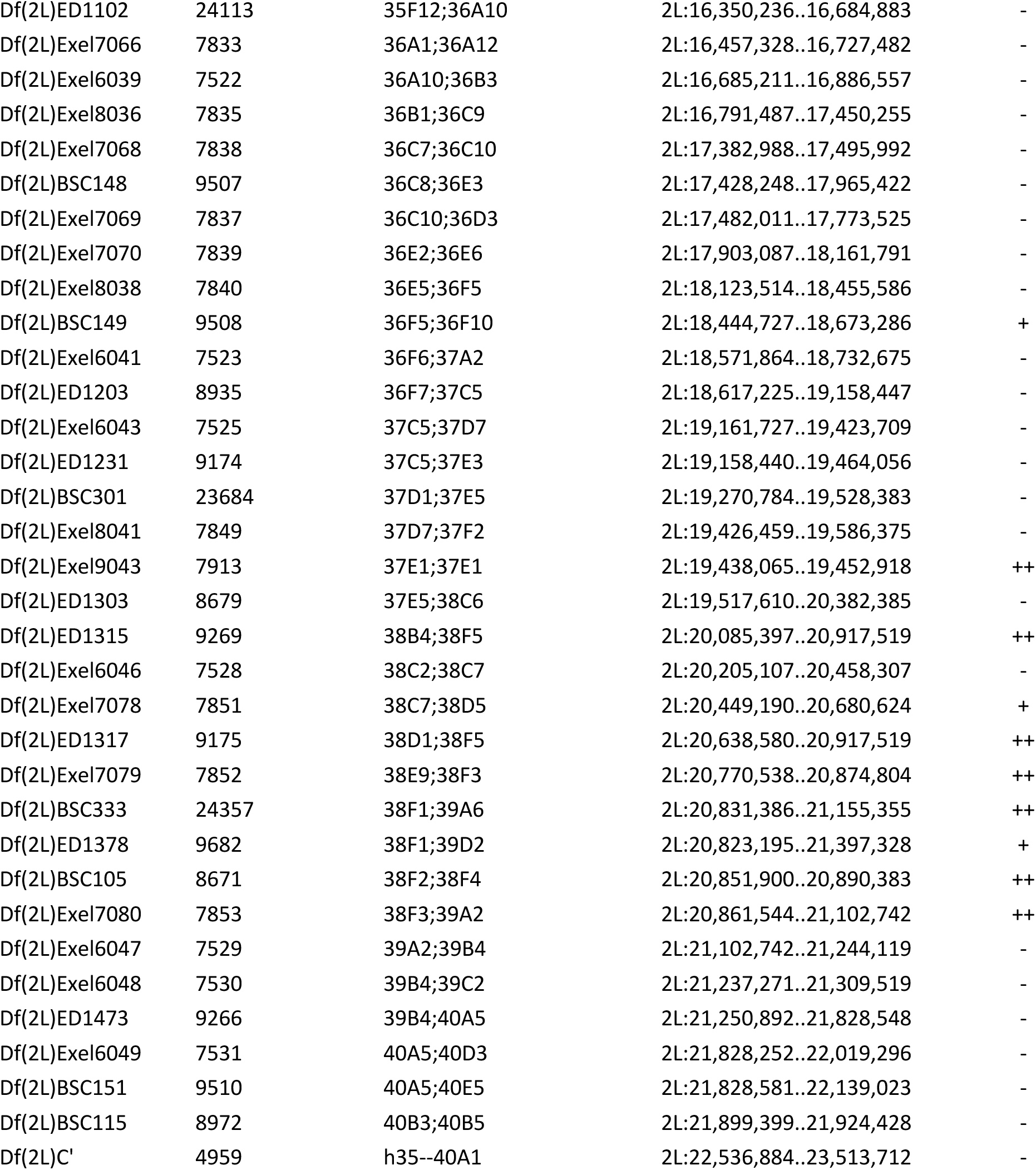
Chromosome 2L deficiencies screened for dominant suppression or enhancement of SigD expression. ^a^ Bloomington Drosophila Stock Center ^b^ as reported by FlyBase Release 6 ^c^ “-” No interaction; “+” Equivocal or weak suppressor; “++” Suppressor; “E” Enhancer. All enhancing deficiencies were assigned a common label rather than differentiating between relatively weak and strong enhancement (as was done with suppressing deficiencies) because of the relative rarity of deficiencies that showed phenotypic enhancement

**Supplementary Table 1b.**
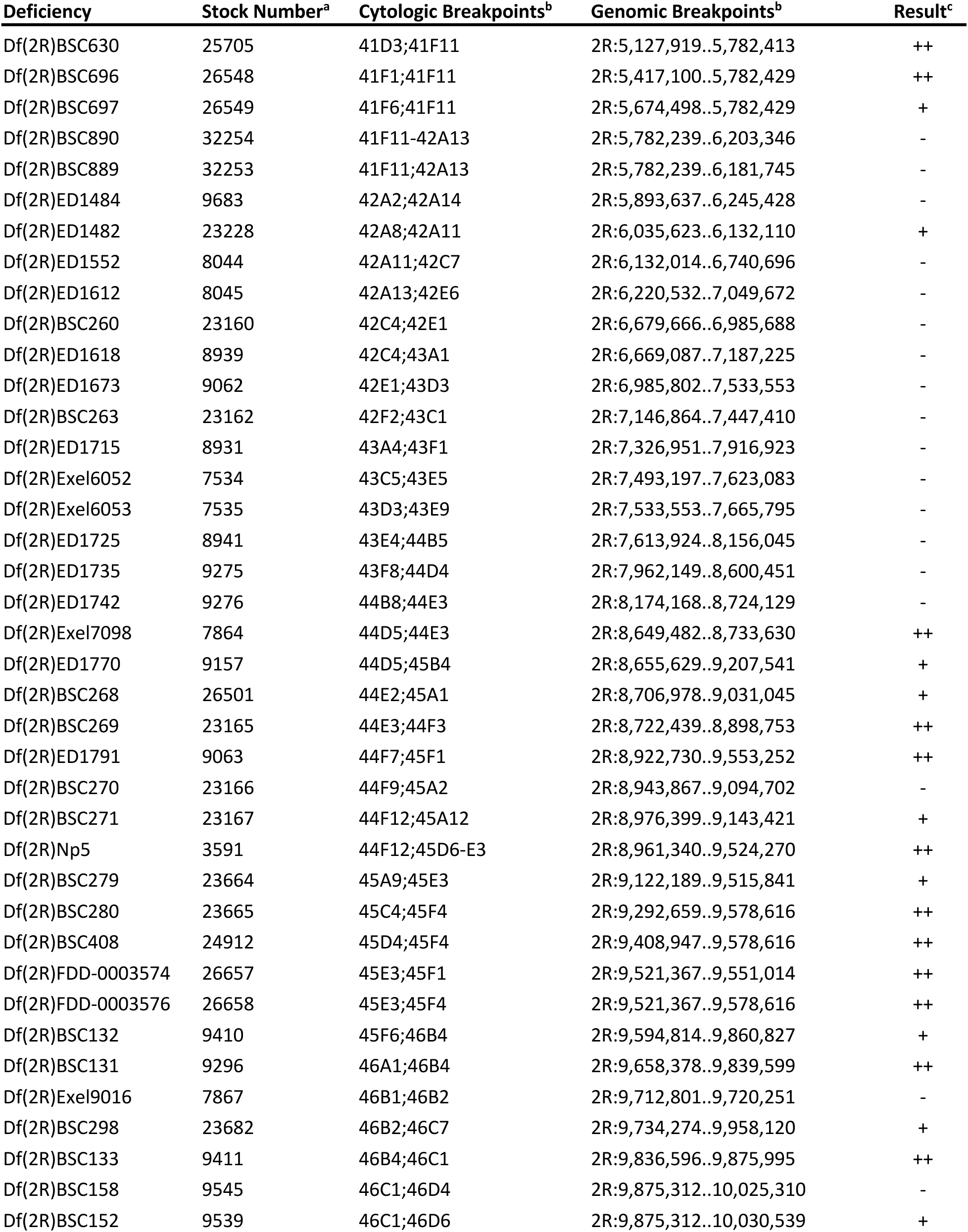
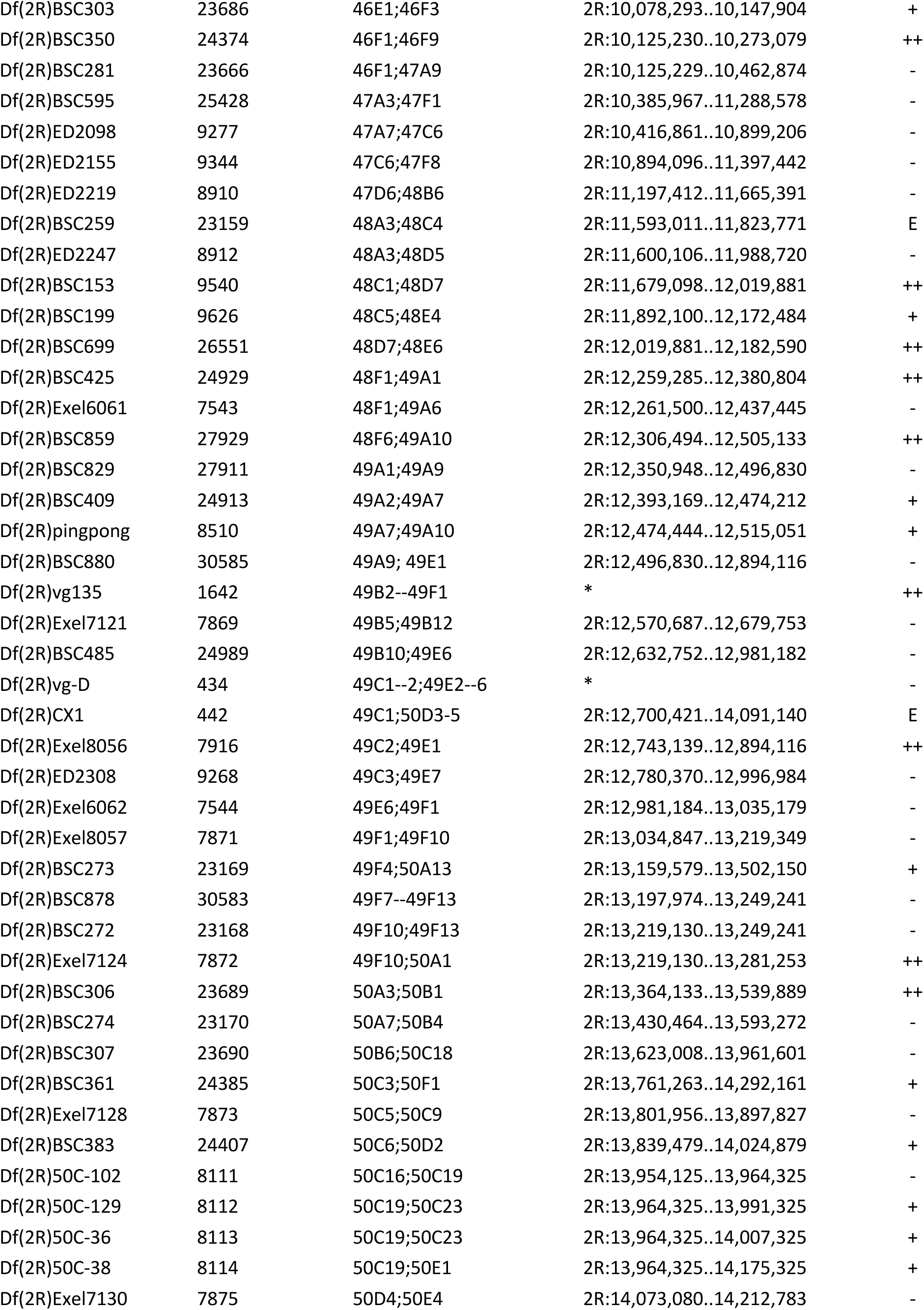
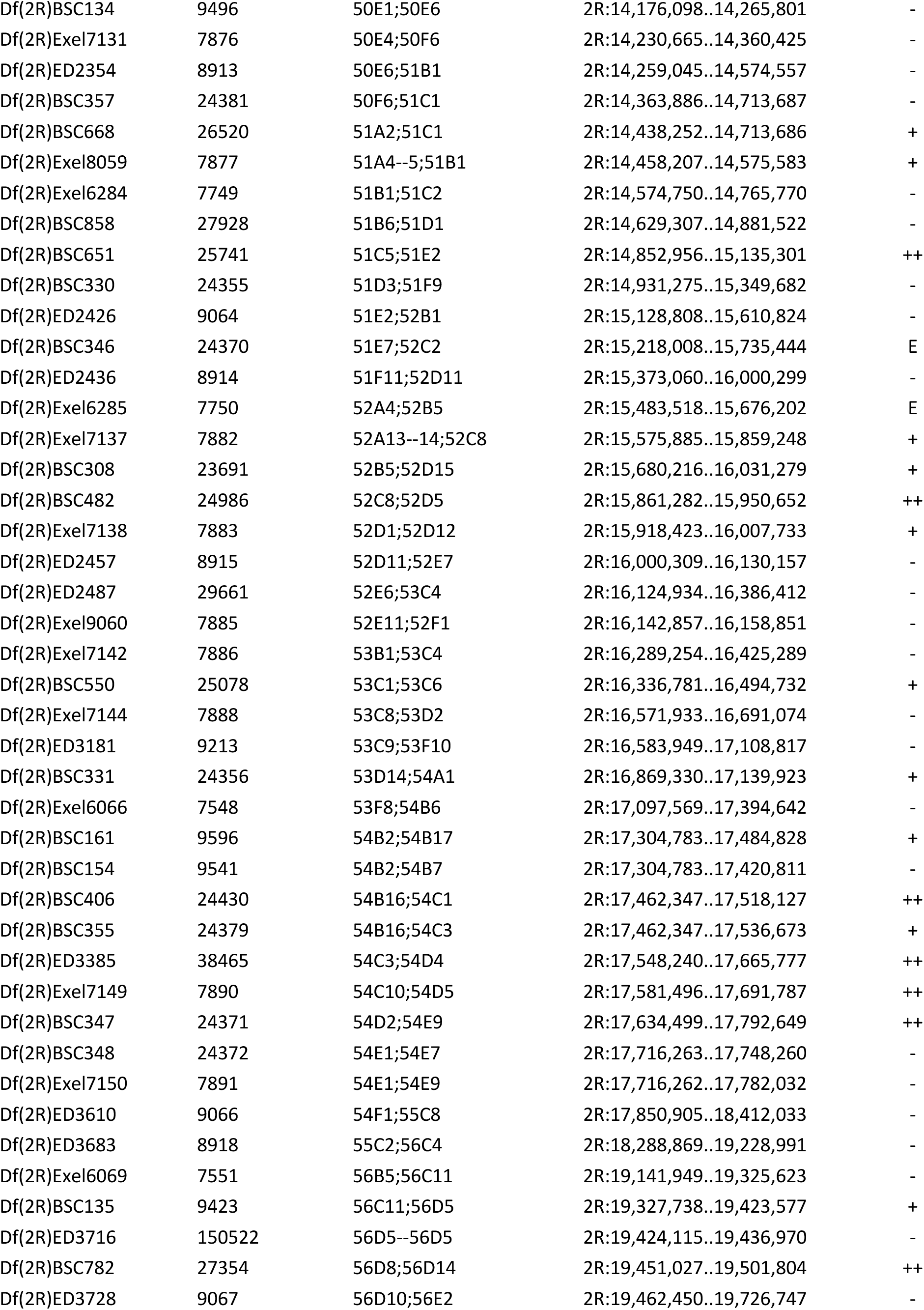
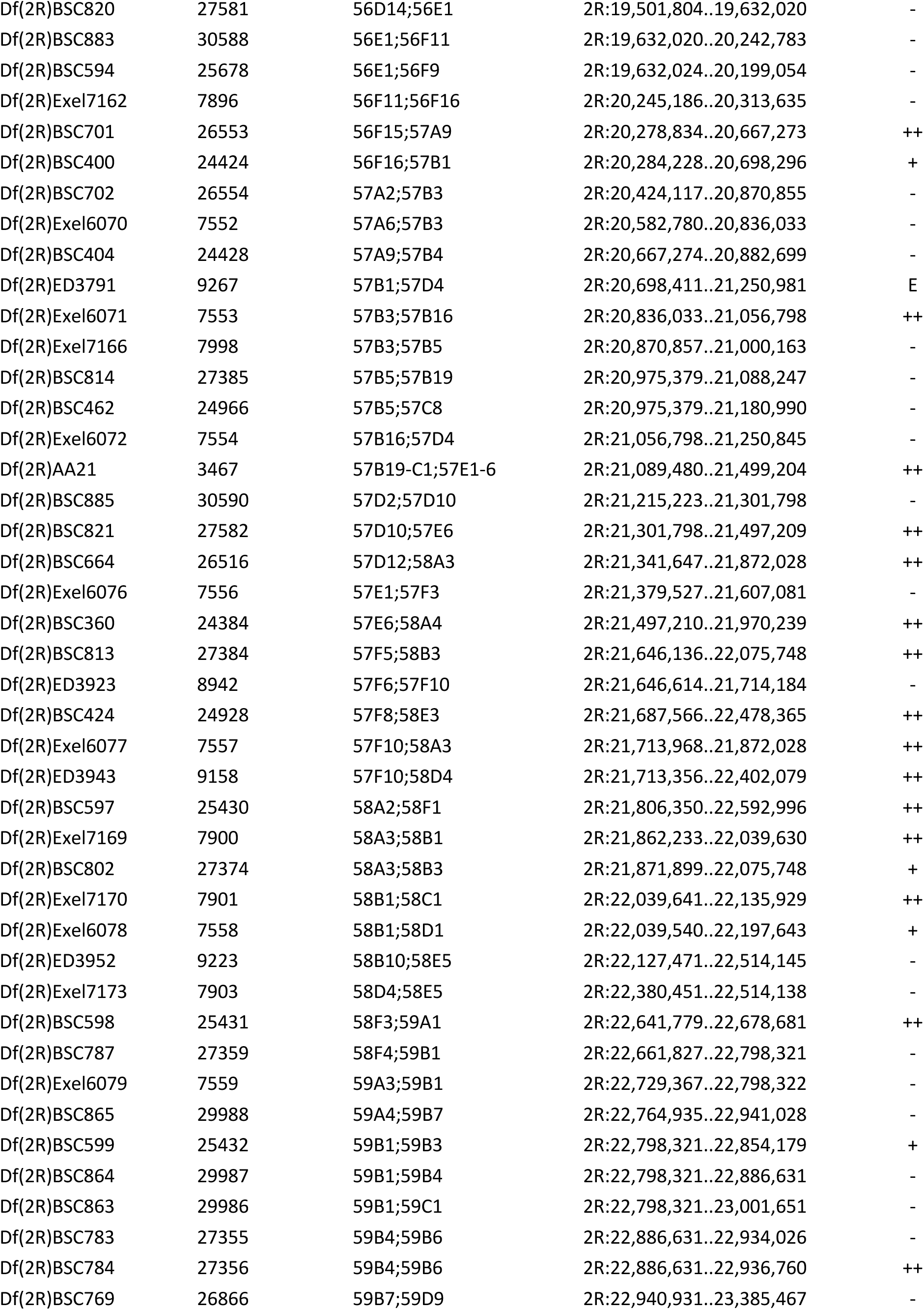
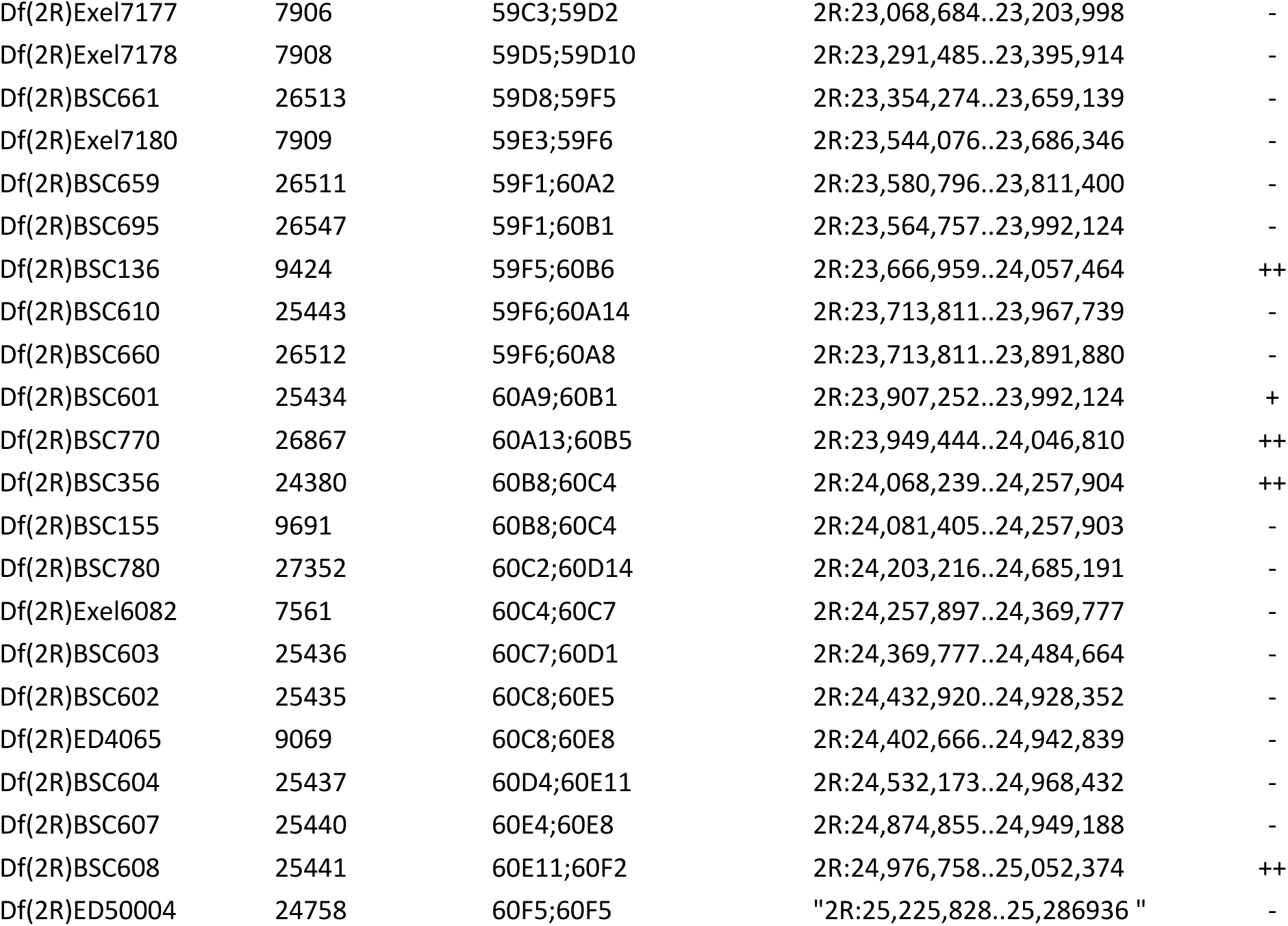
Chromosome 2R deficiencies screened for dominant suppression or enhancement of SigD expression. ^a^ Bloomington Drosophila Stock Center ^b^ as reported by FlyBase Release 6 ^c^ “-” no interaction; “+” equivocal or weak suppressor; “++” suppressor; “E” enhancer

**Supplementary Table 1c.**
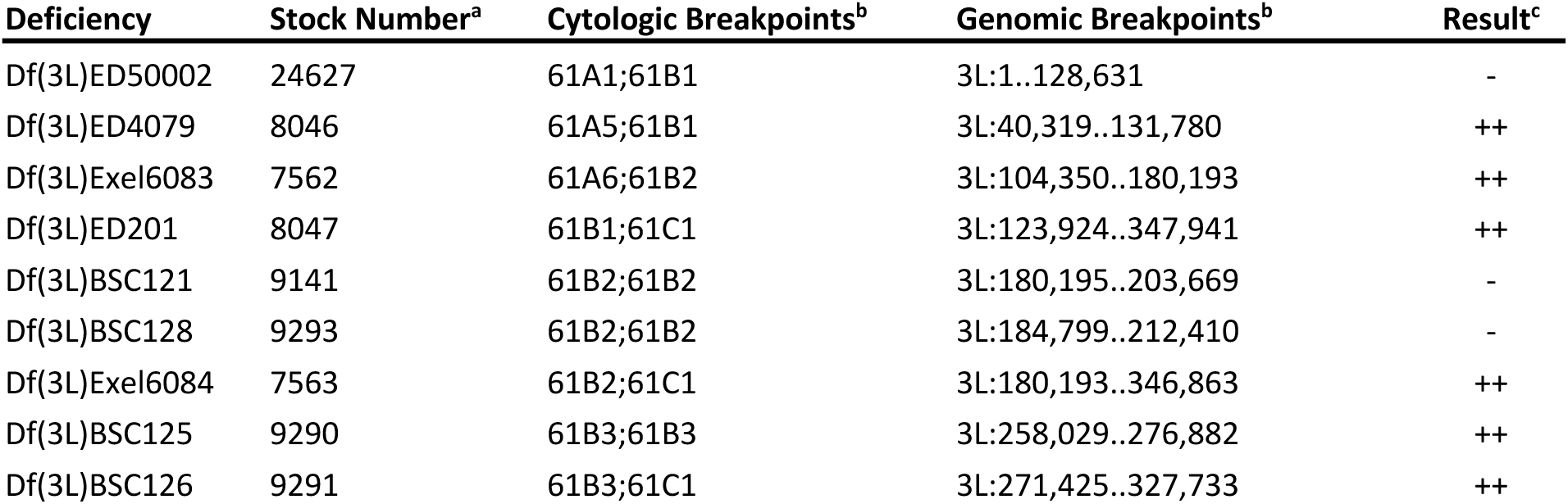
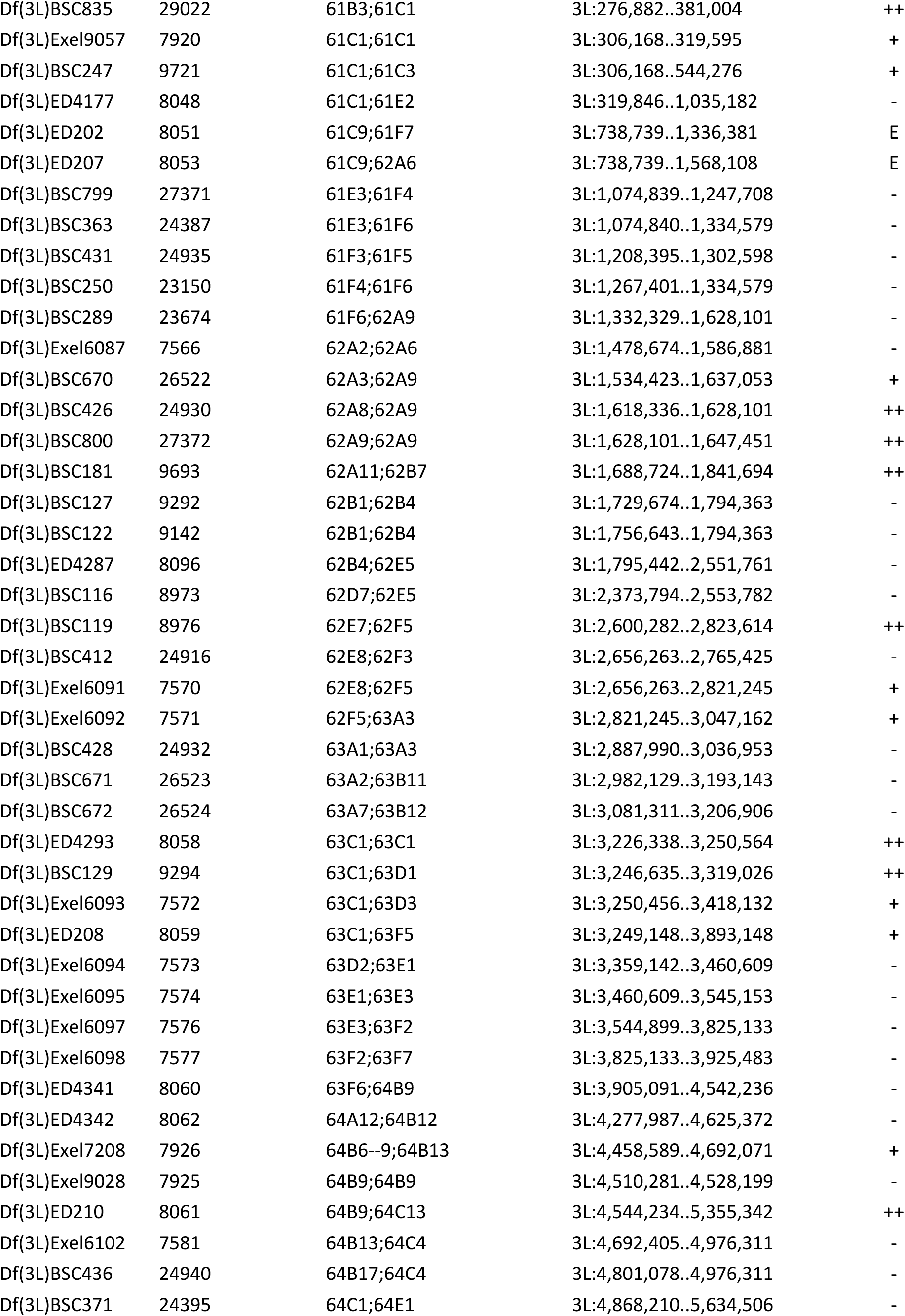
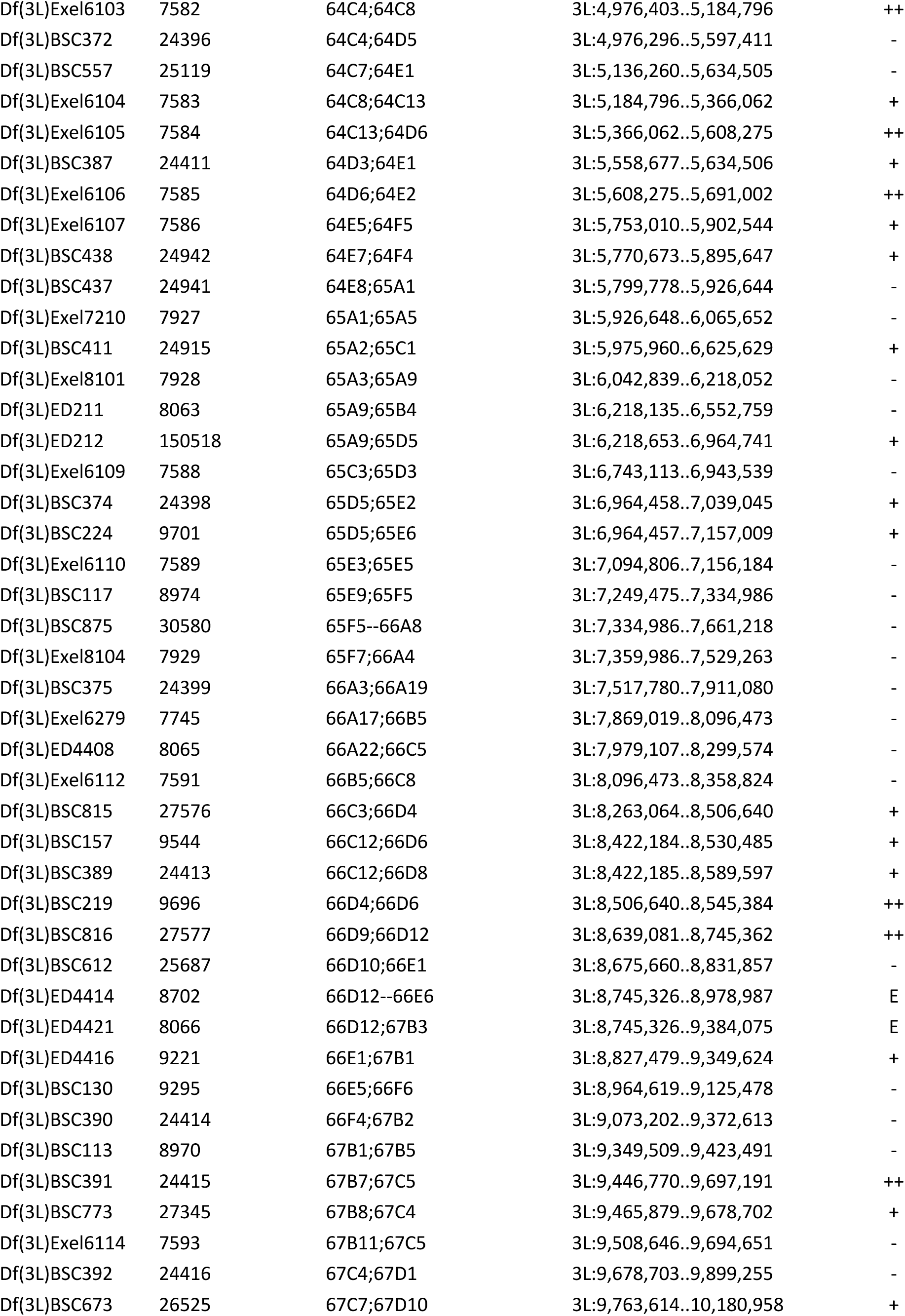
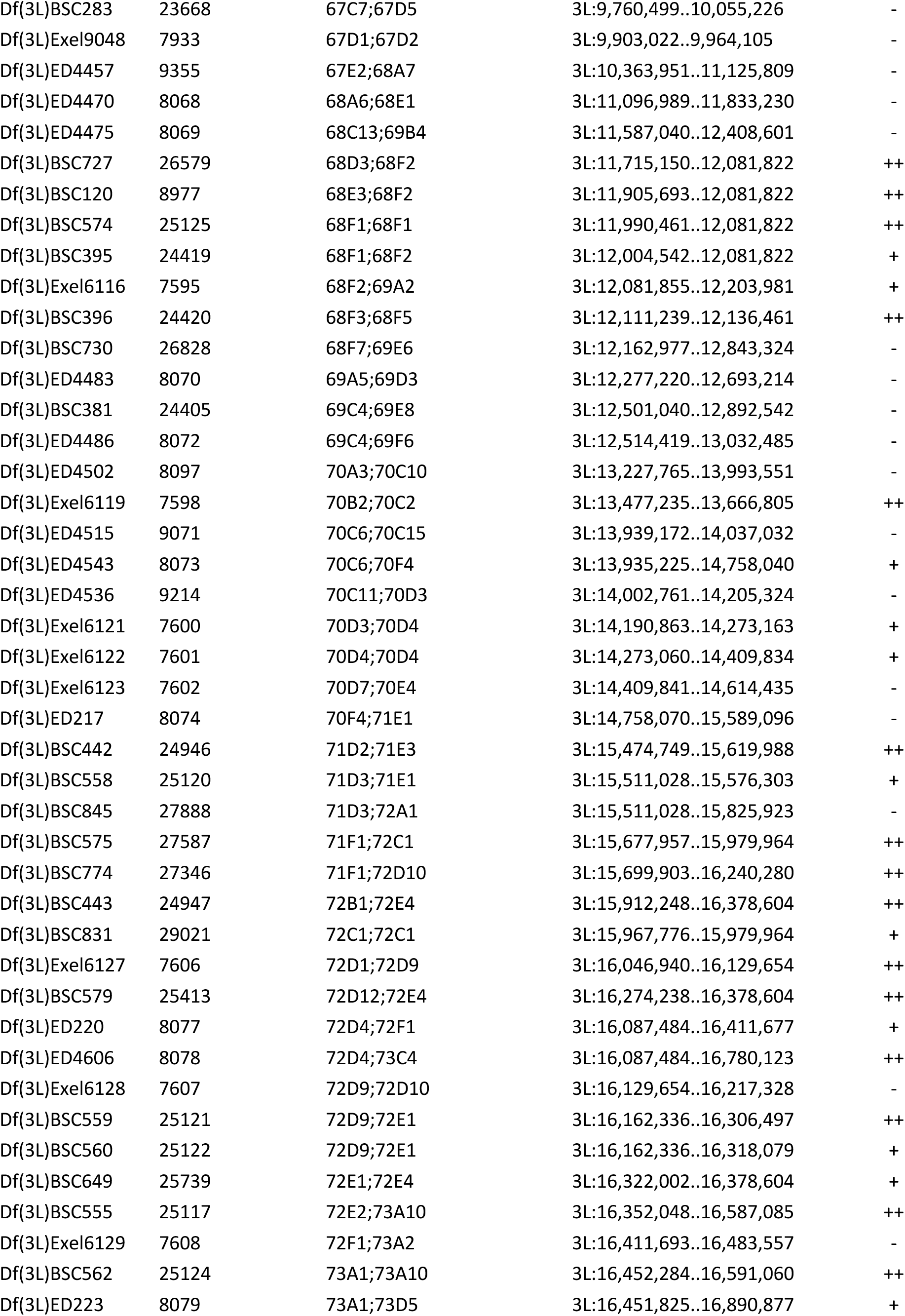
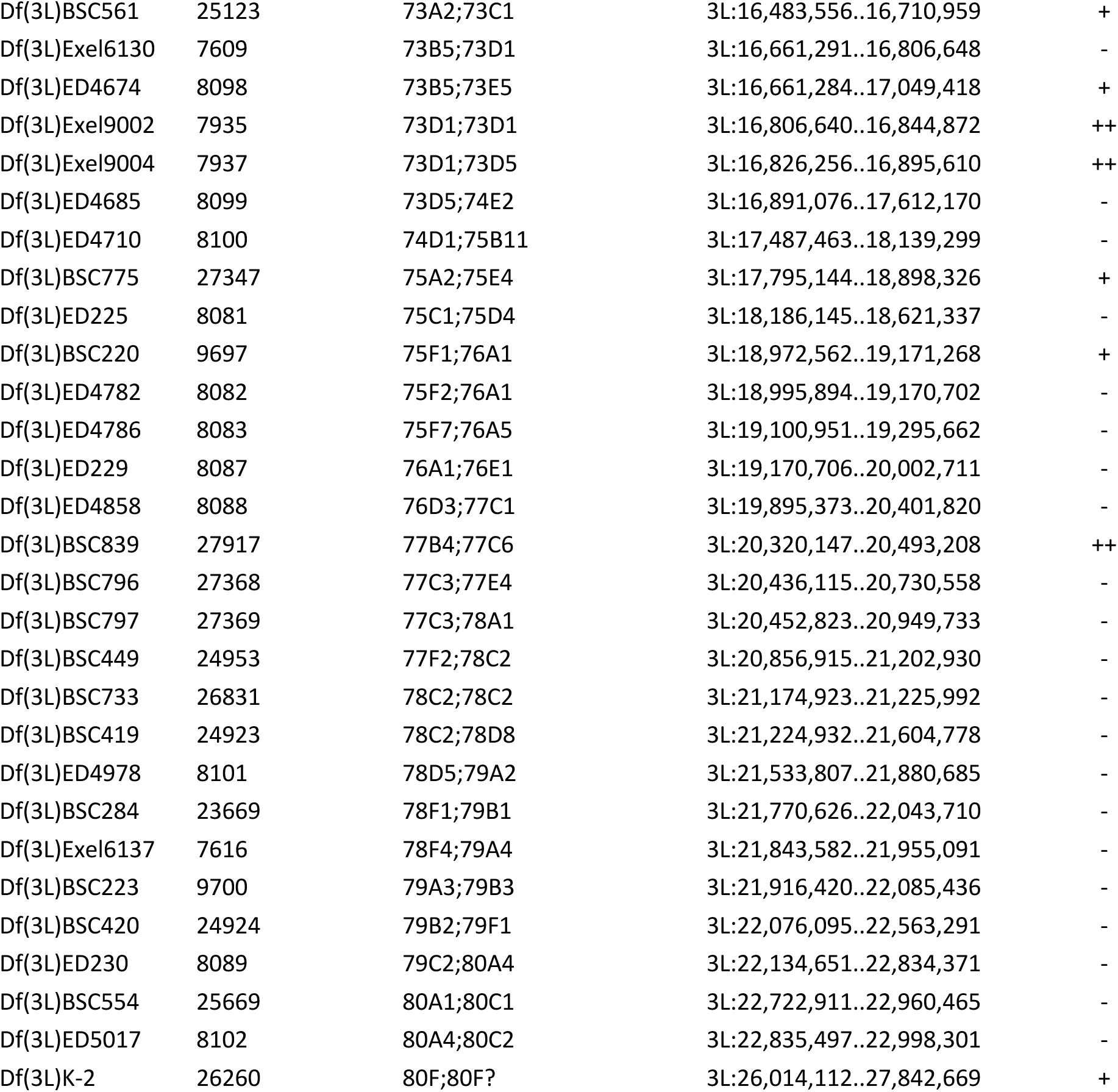
Chromosome 3L deficiencies screened for dominant suppression or enhancement of SigD expression. ^a^ Bloomington Drosophila Stock Center ^b^ as reported by FlyBase Release 6 ^c^ “-” no interaction; “+” equivocal or weak suppressor; “++” suppressor; “E” enhancer

**Supplementary Table 1d.**
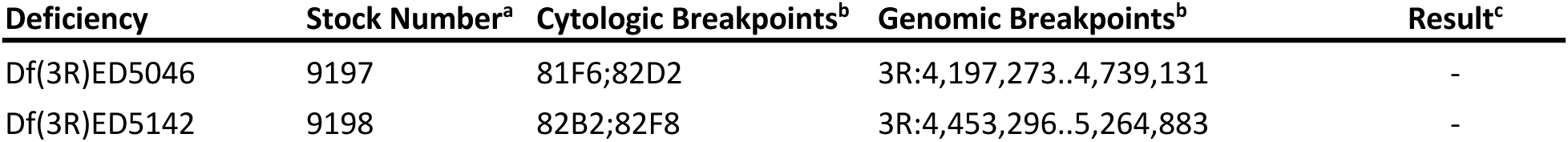
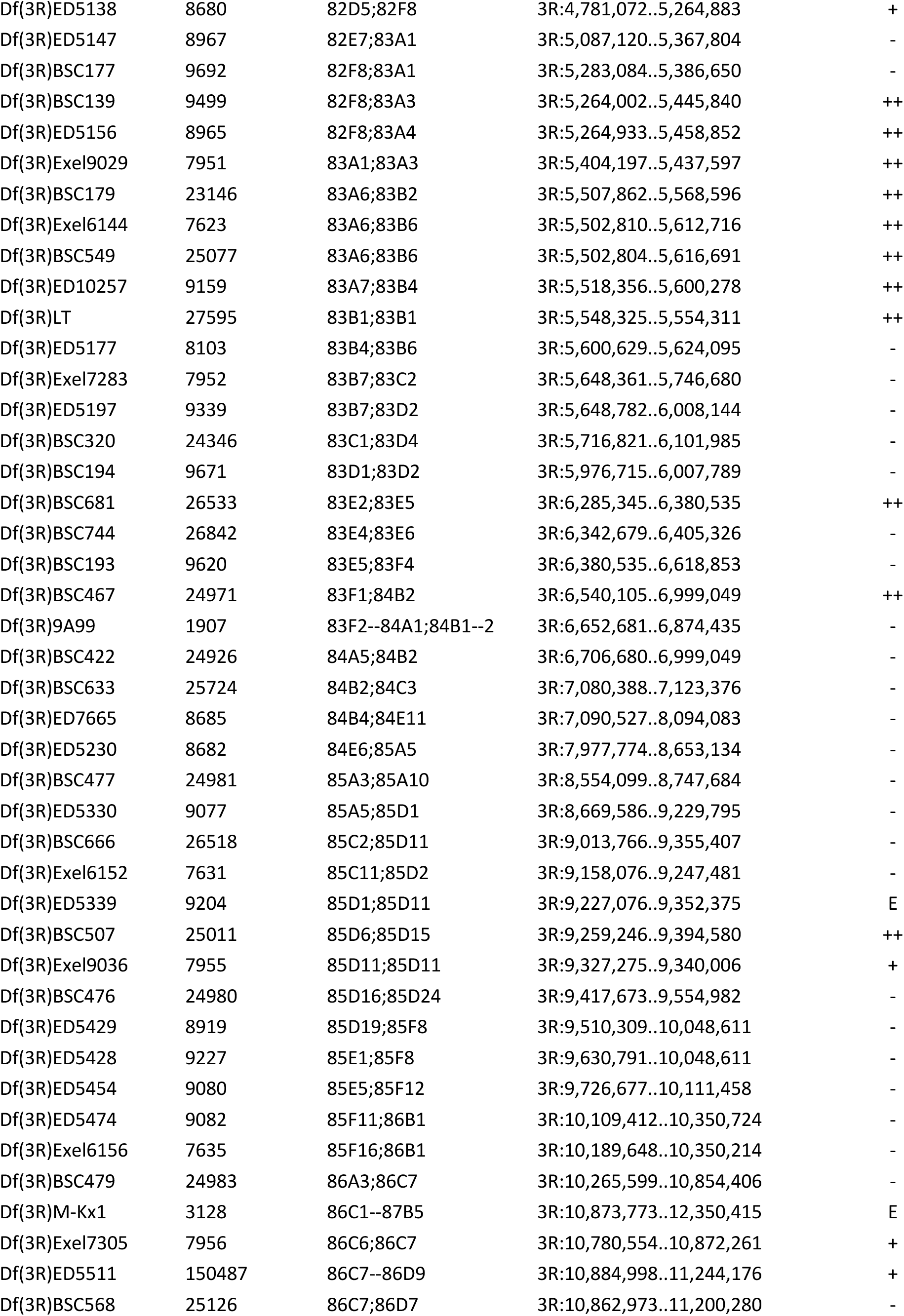
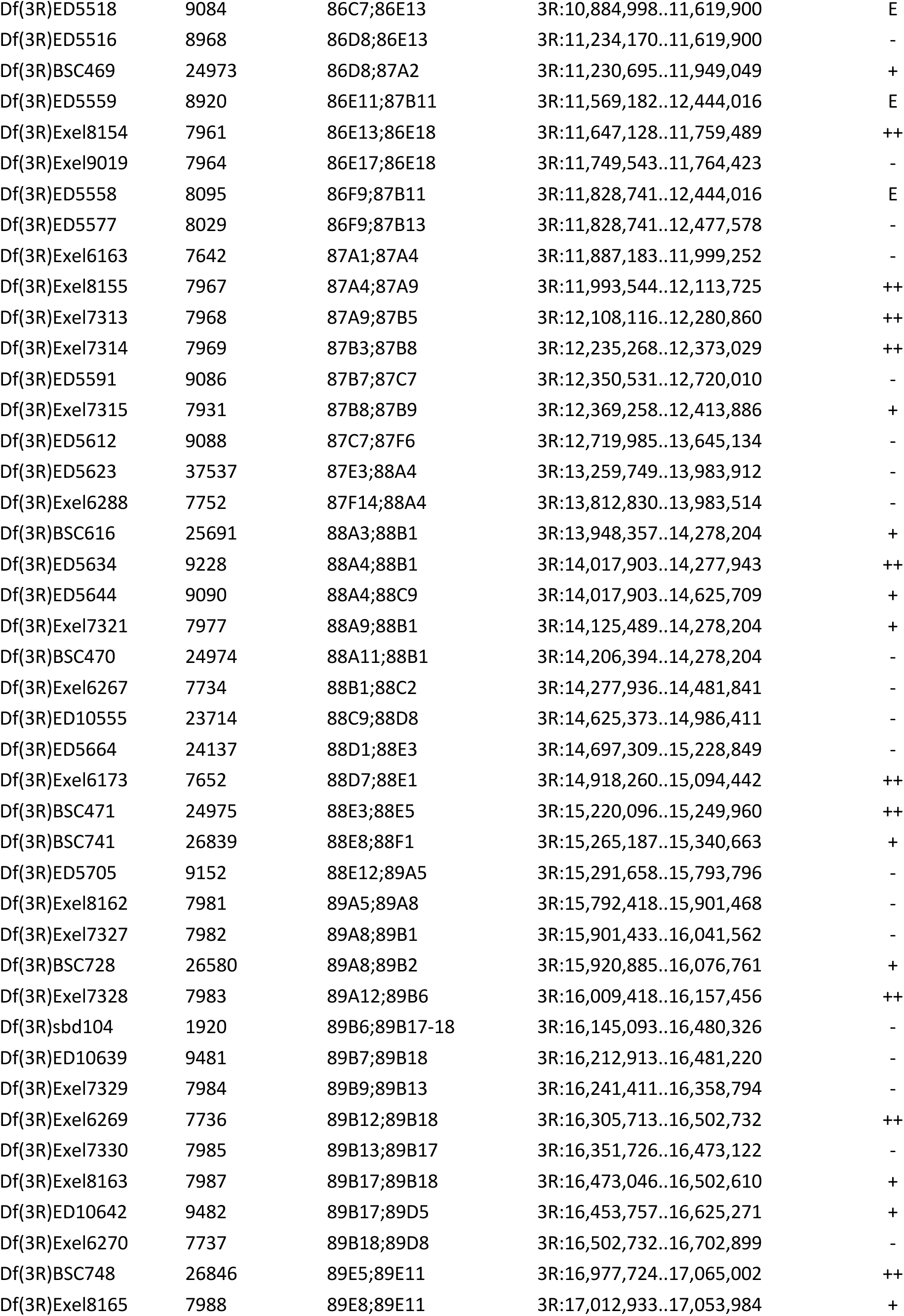
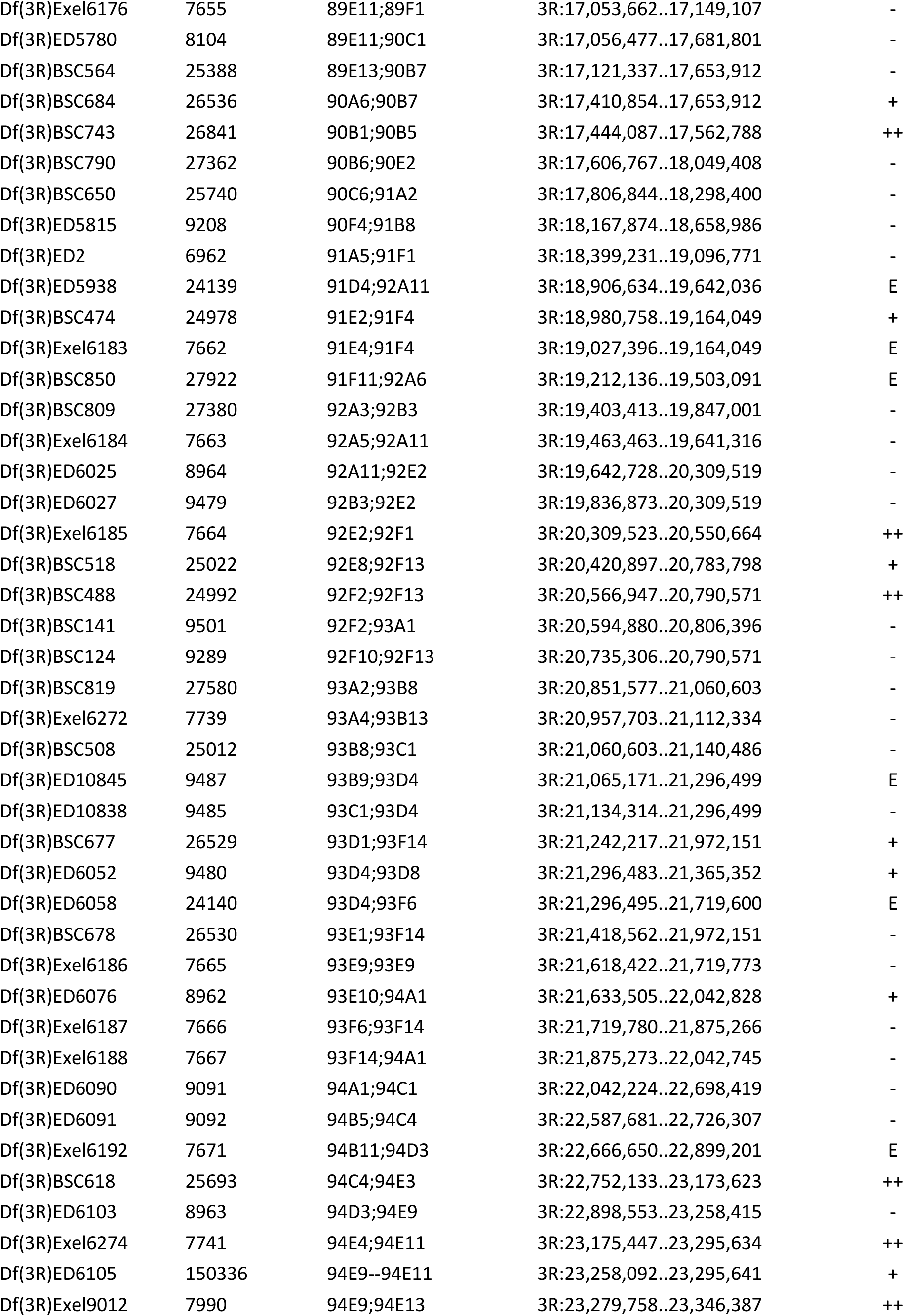
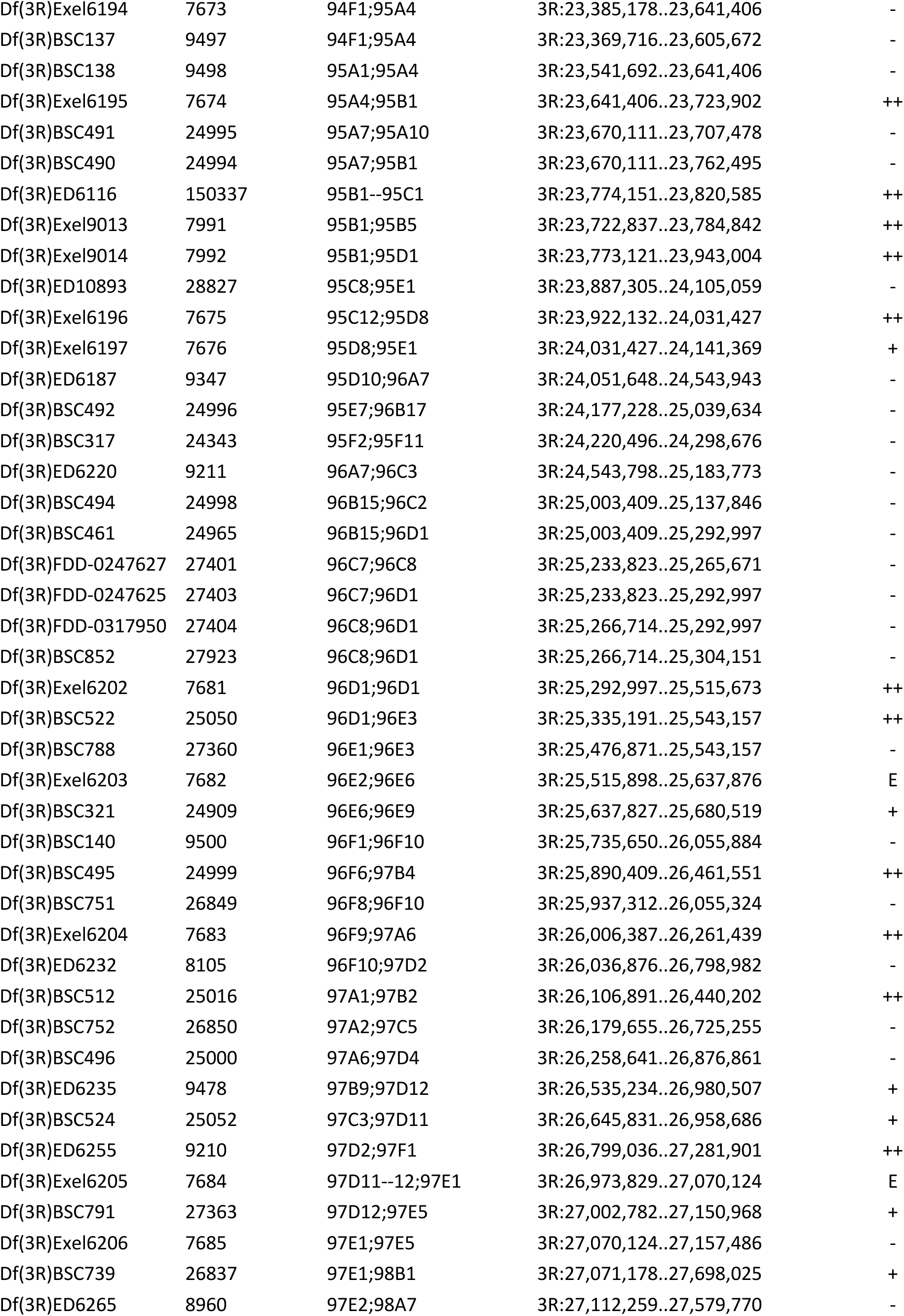
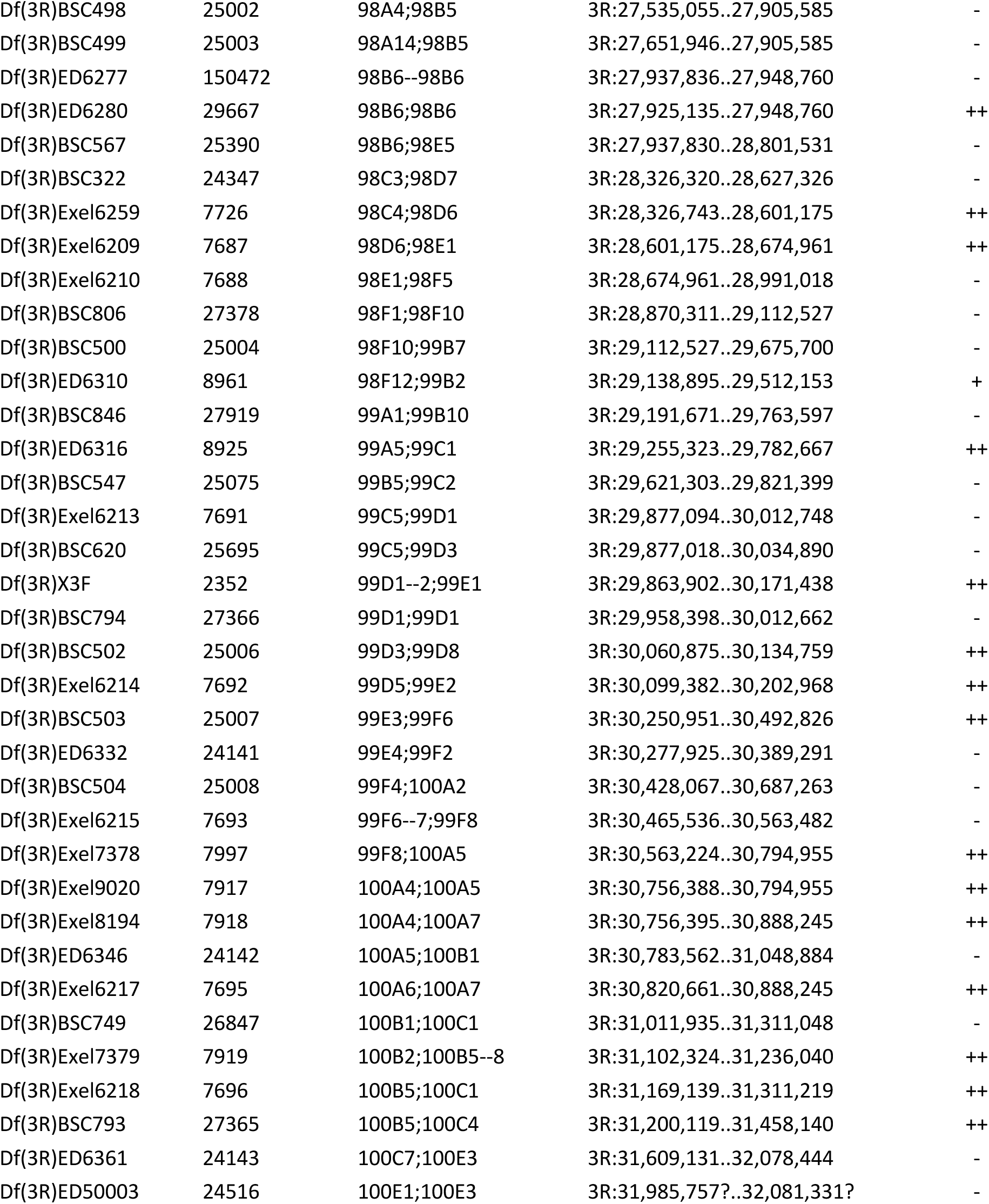
Chromosome 3R deficiencies screened for dominant suppression or enhancement of SigD expression. ^a^ Bloomington Drosophila Stock Center ^b^ as reported by FlyBase Release 6 ^c^ “-” no interaction; “+” equivocal or weak suppressor; “++” suppressor; “E” enhancer

**Supplementary Table 2.**
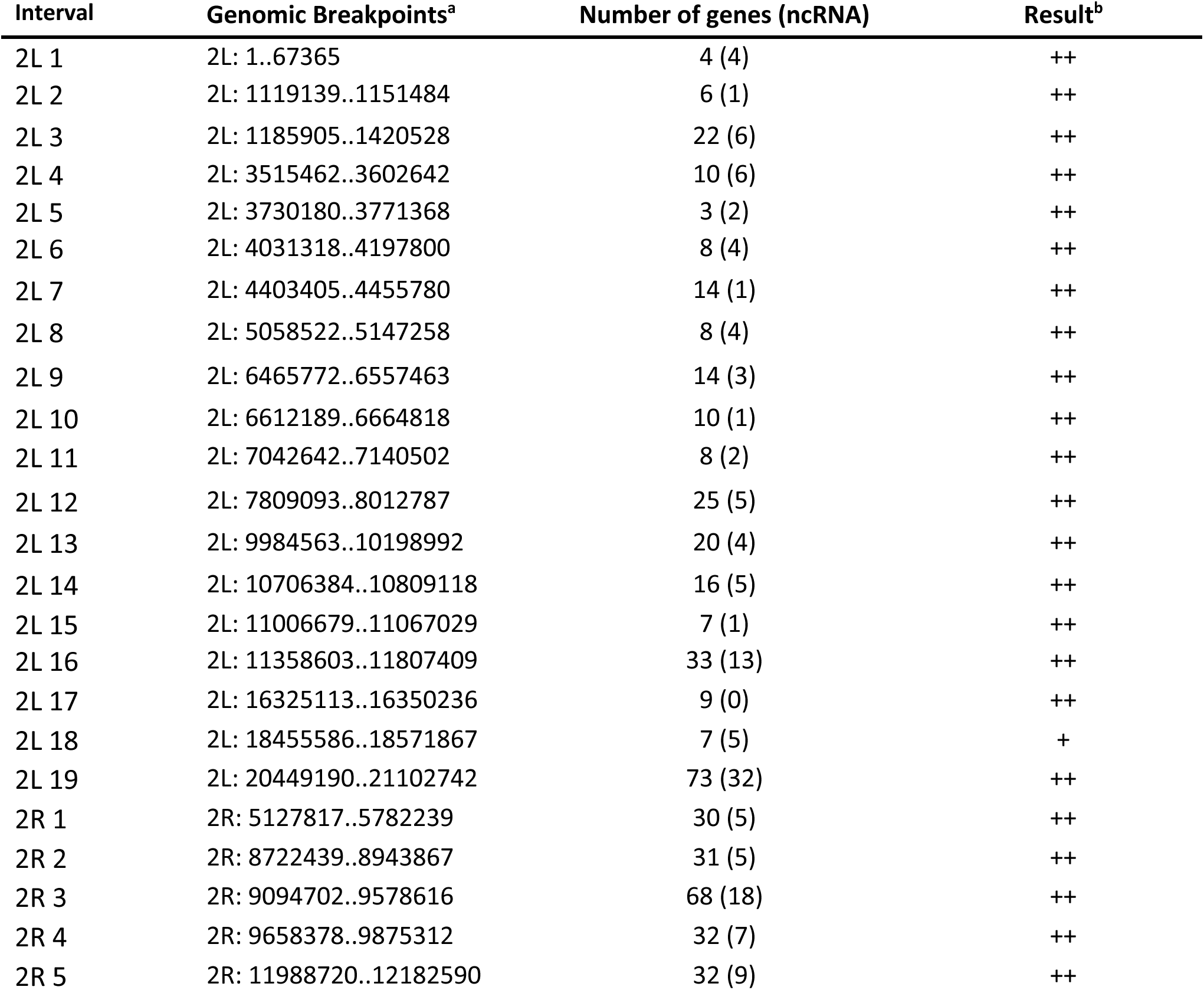
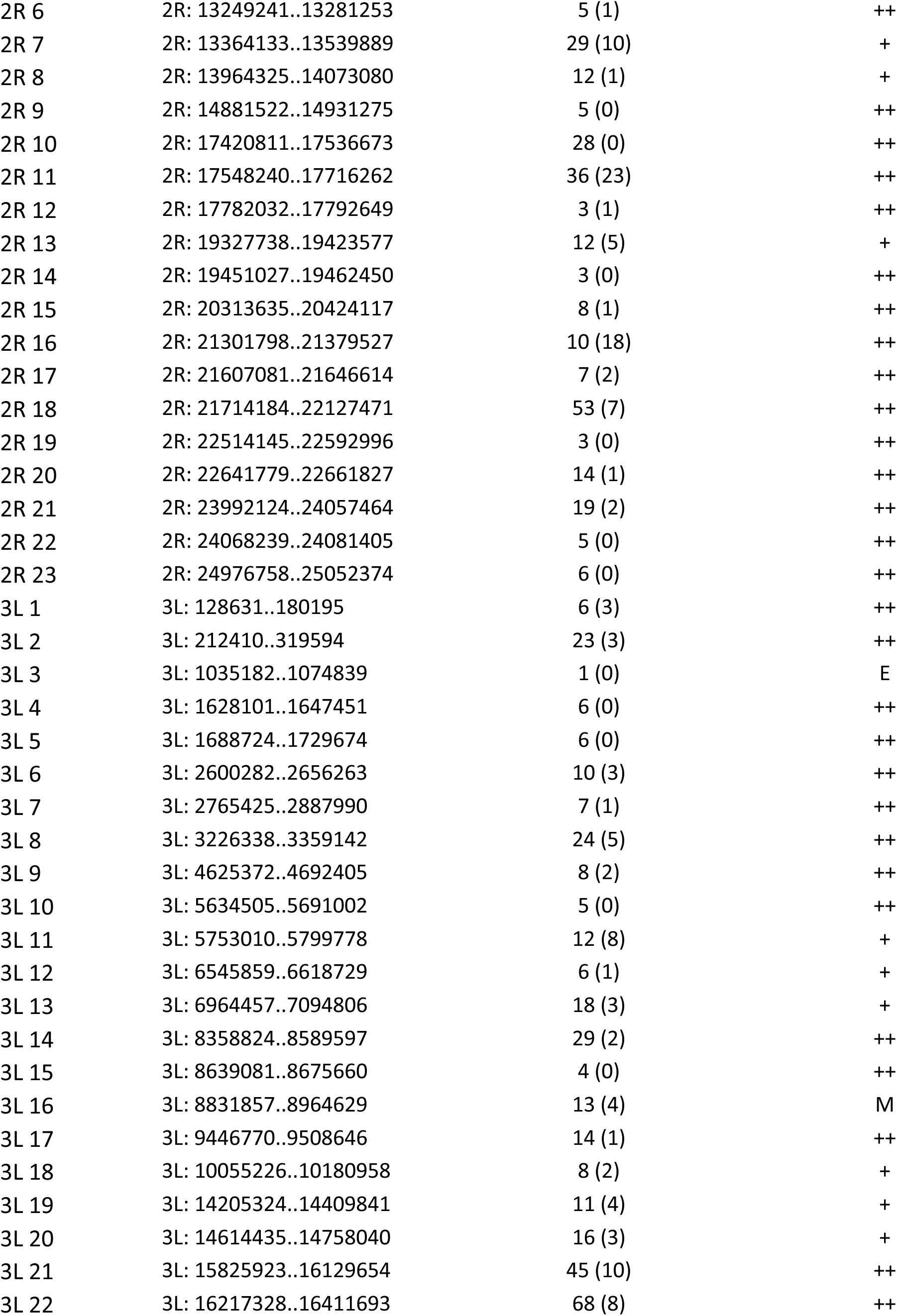
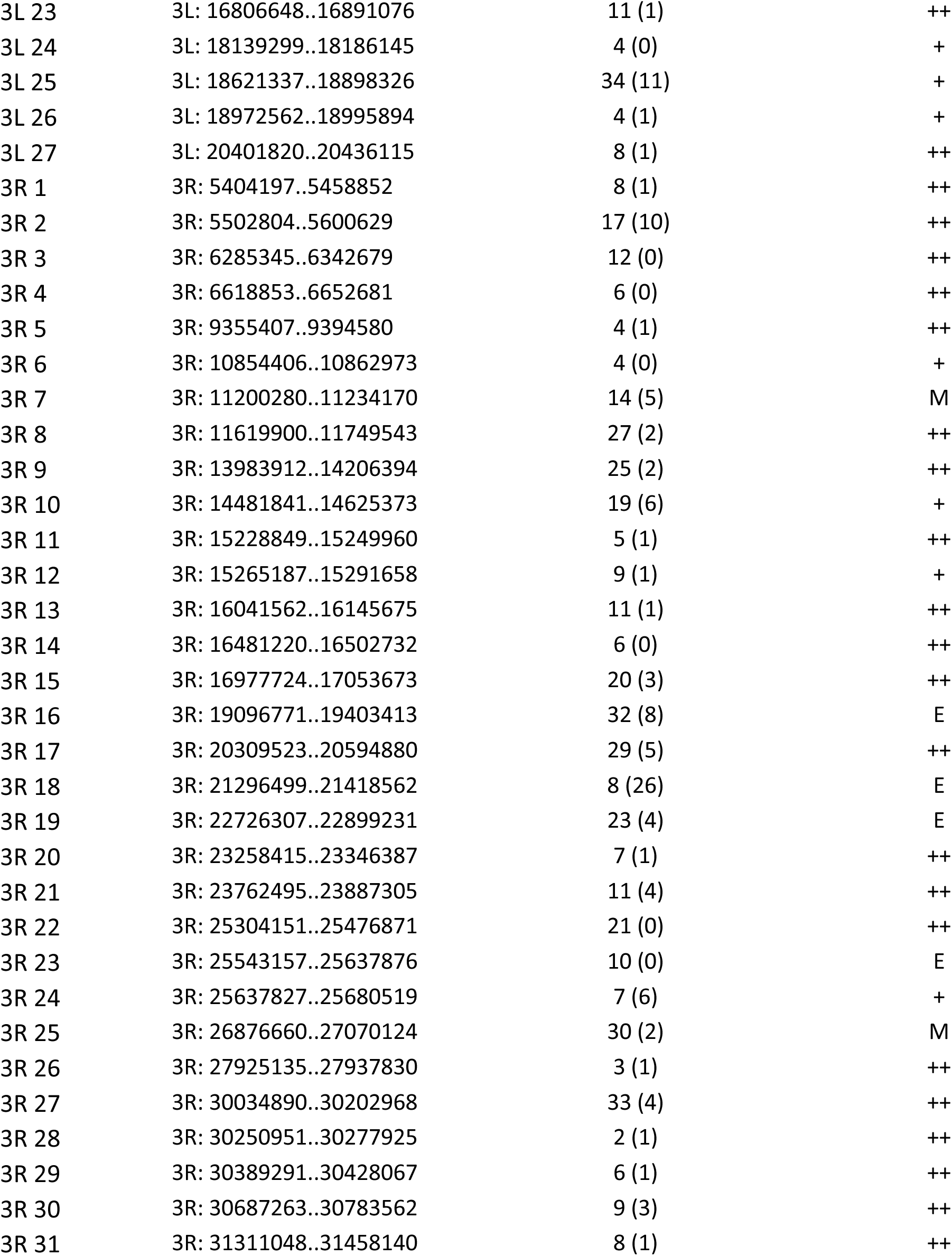
Interacting genomic intervals derived from screening chromosomal deficiencies for dominant suppression or enhancement of SigD expression. ^a^ interval breakpoints correspond to the breakpoints of chromosomal deficiencies that define the interval, as reported by the FlyBase Consortium (2020) ^b^ SUPPRESSOR (++): the interval is defined by a single or overlapping suppressor deficiencies (as described in Supp Table 1) WEAK SUPPRESSOR (+): the interval is defined by a single or overlapping weak suppressor deficiencies (as described in Supp Table 1) ENHANCER (E): the interval is defined by one or more enhancer deficiencies (as described in Supp Table 1) MIXED (M): the interval is defined by one or more suppressing deficiencies, overlapped by one or more enhancing deficiencies

**Supplementary Table 3.**
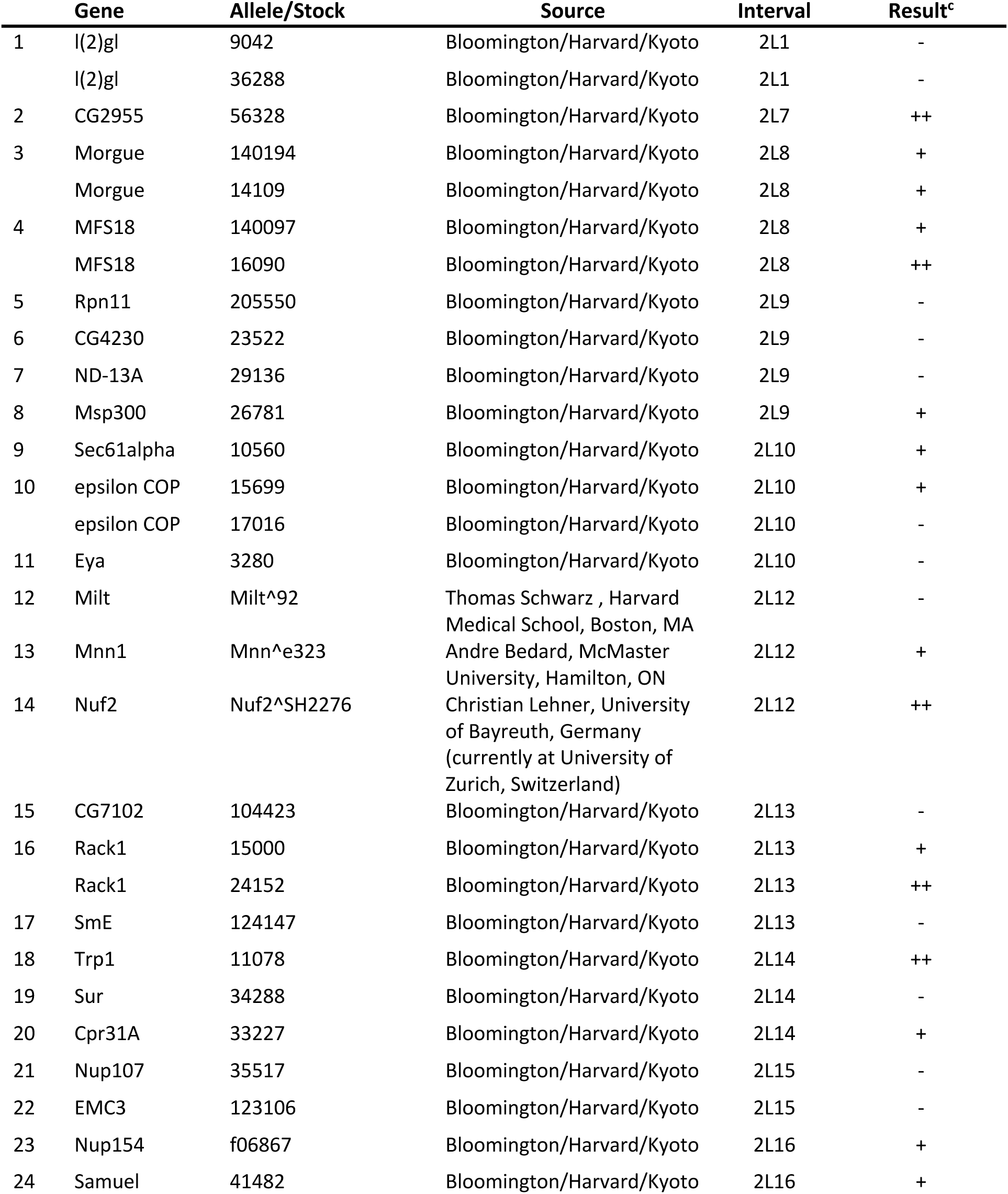
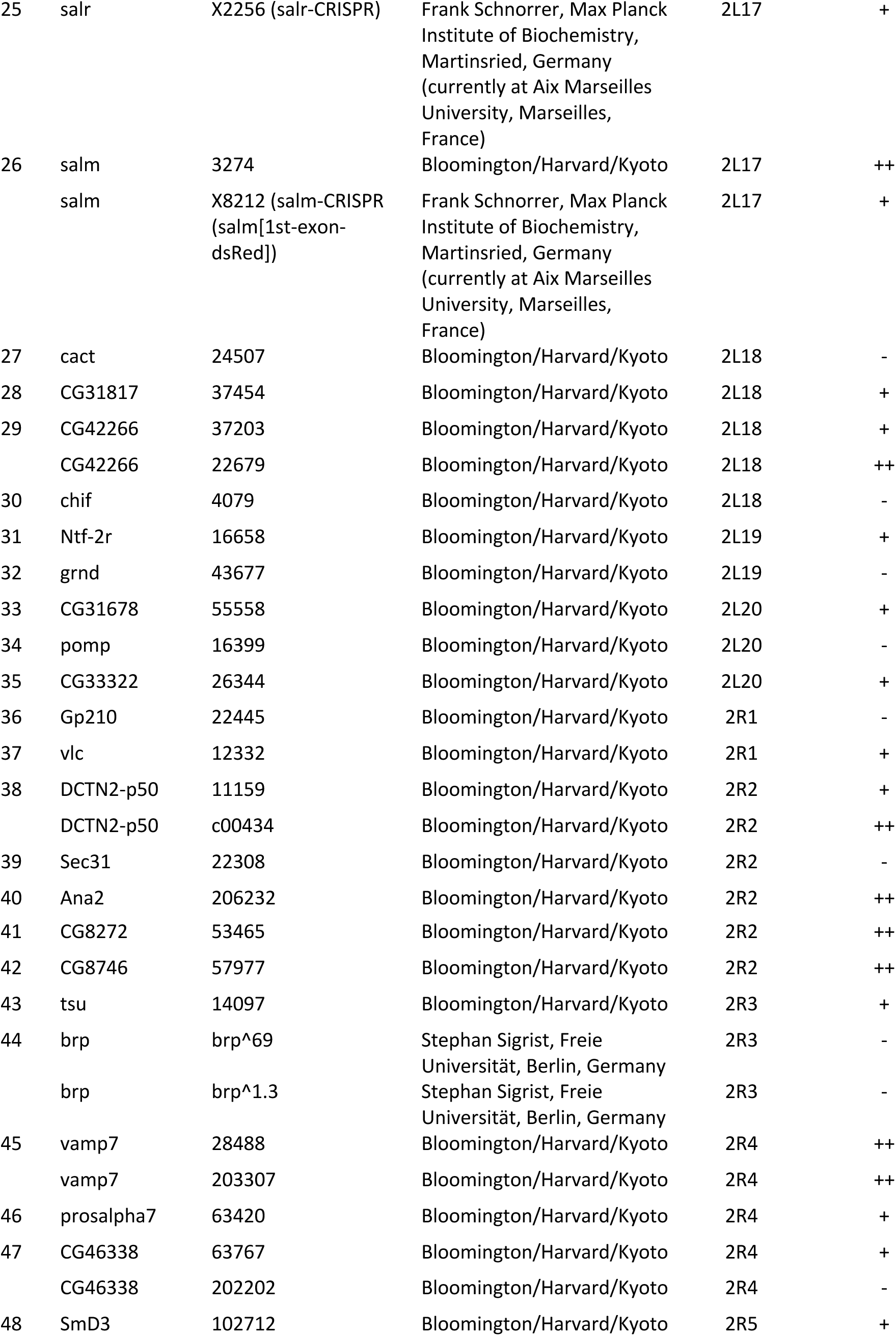
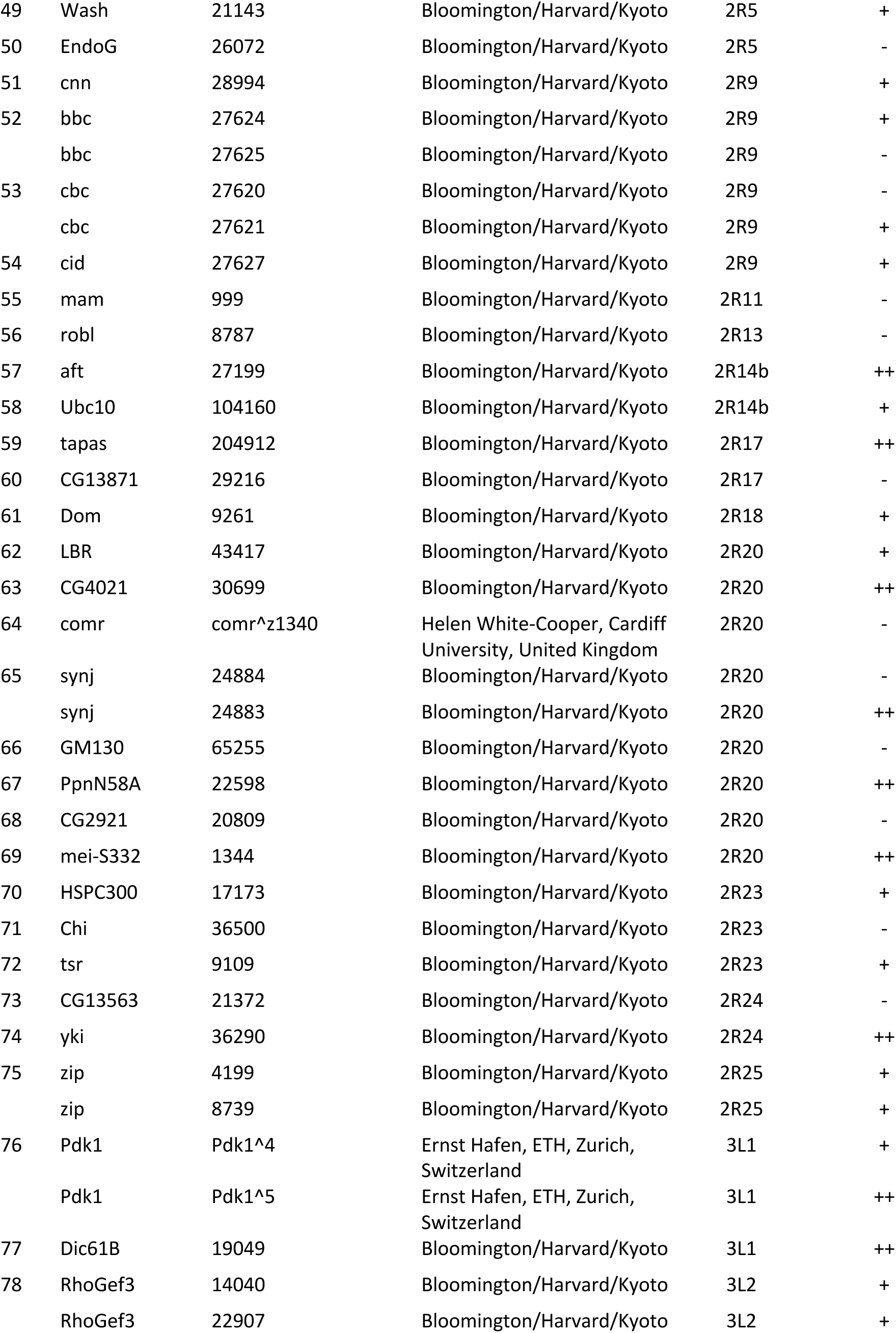
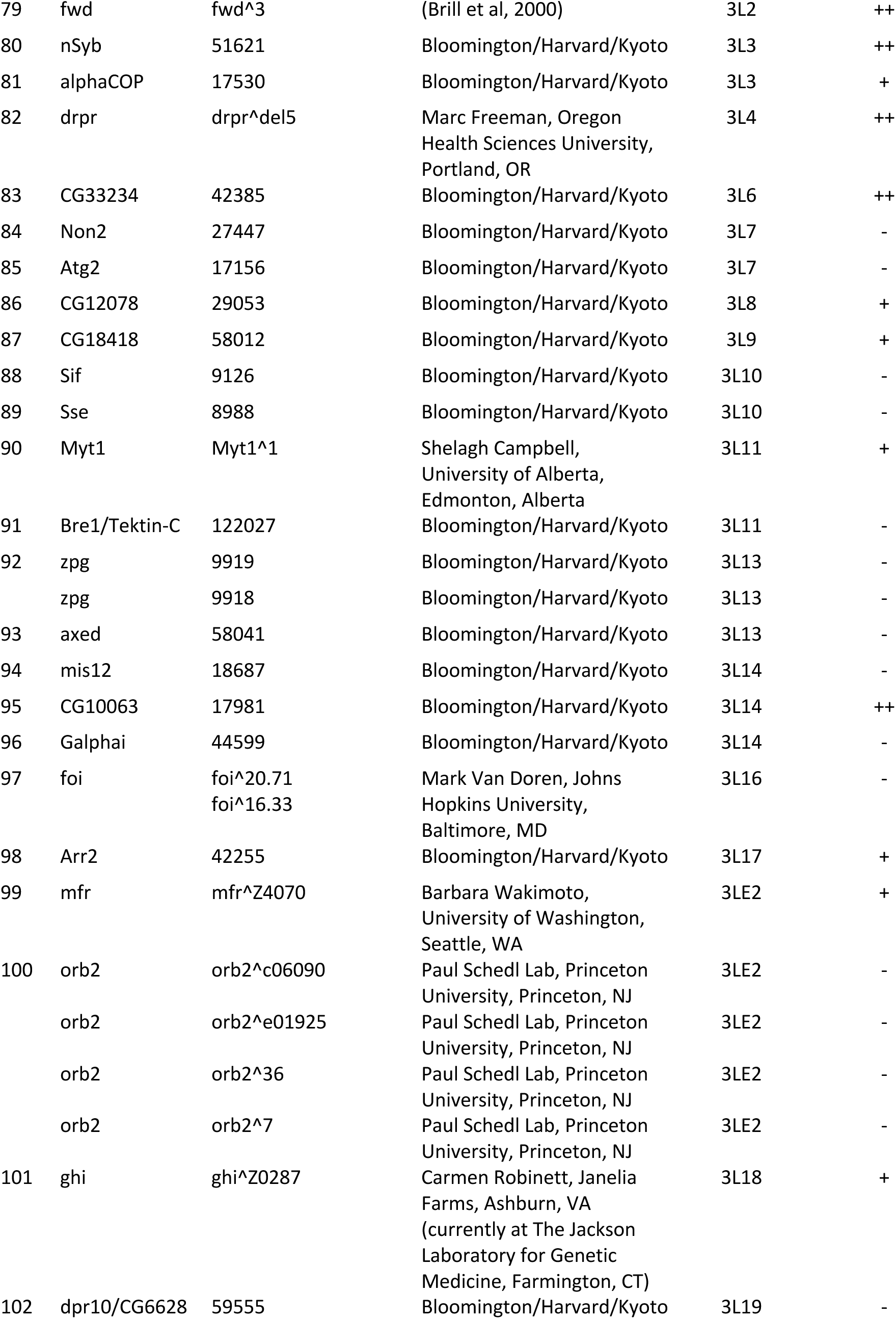
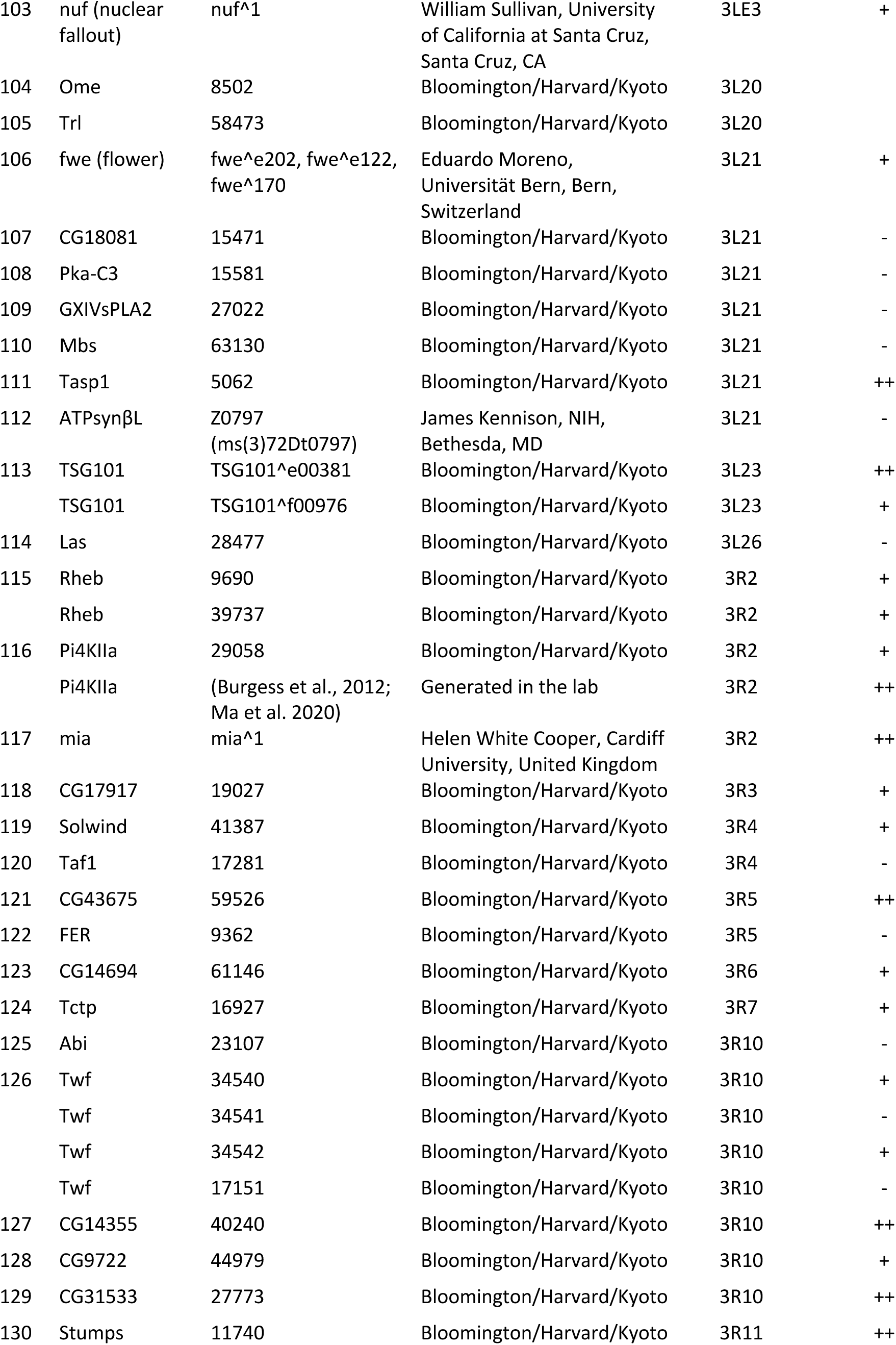
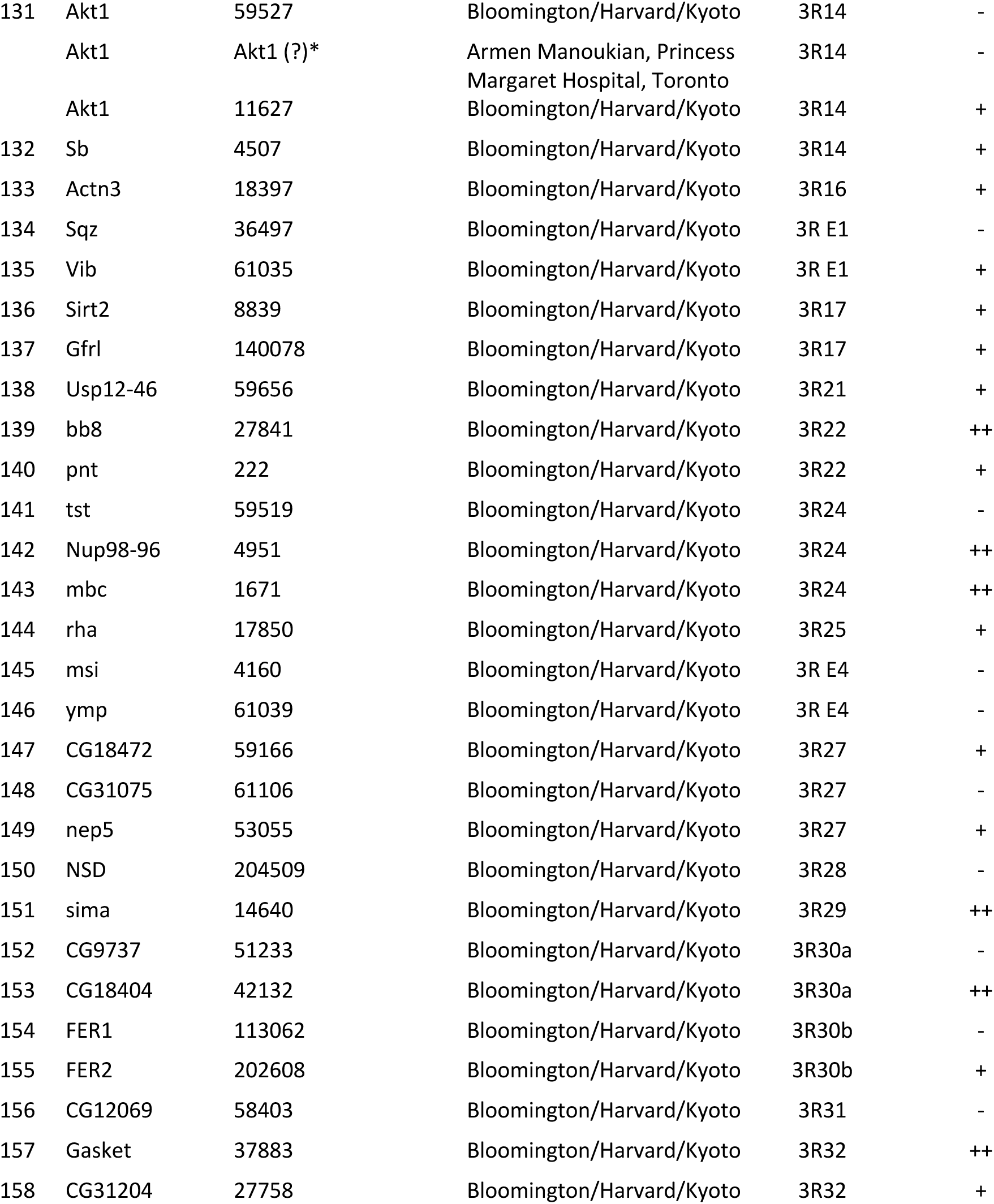
^c^ “-” No interaction; “+” Equivocal or weak suppressor; “++” Suppressor

**Supplementary Table 4.**
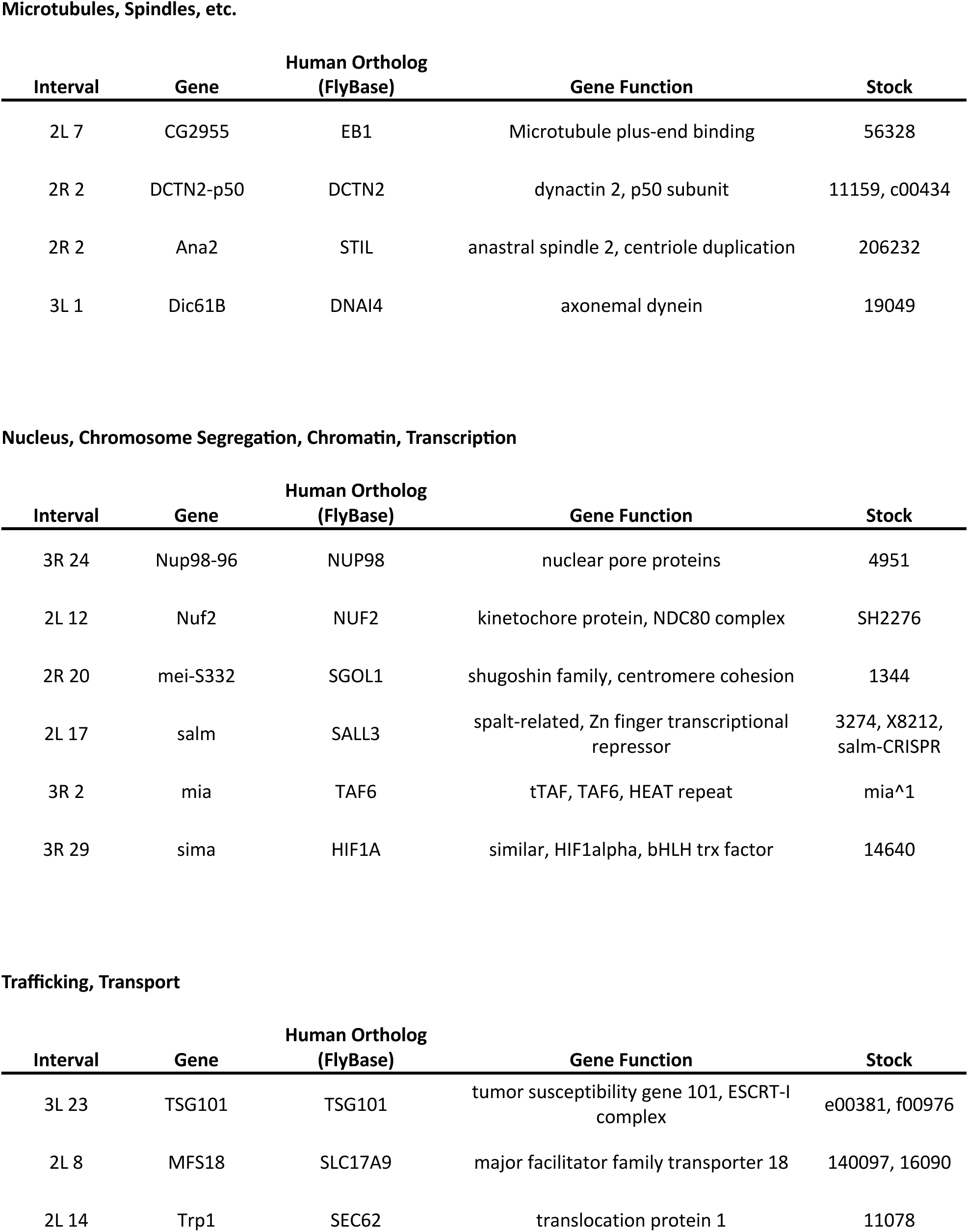
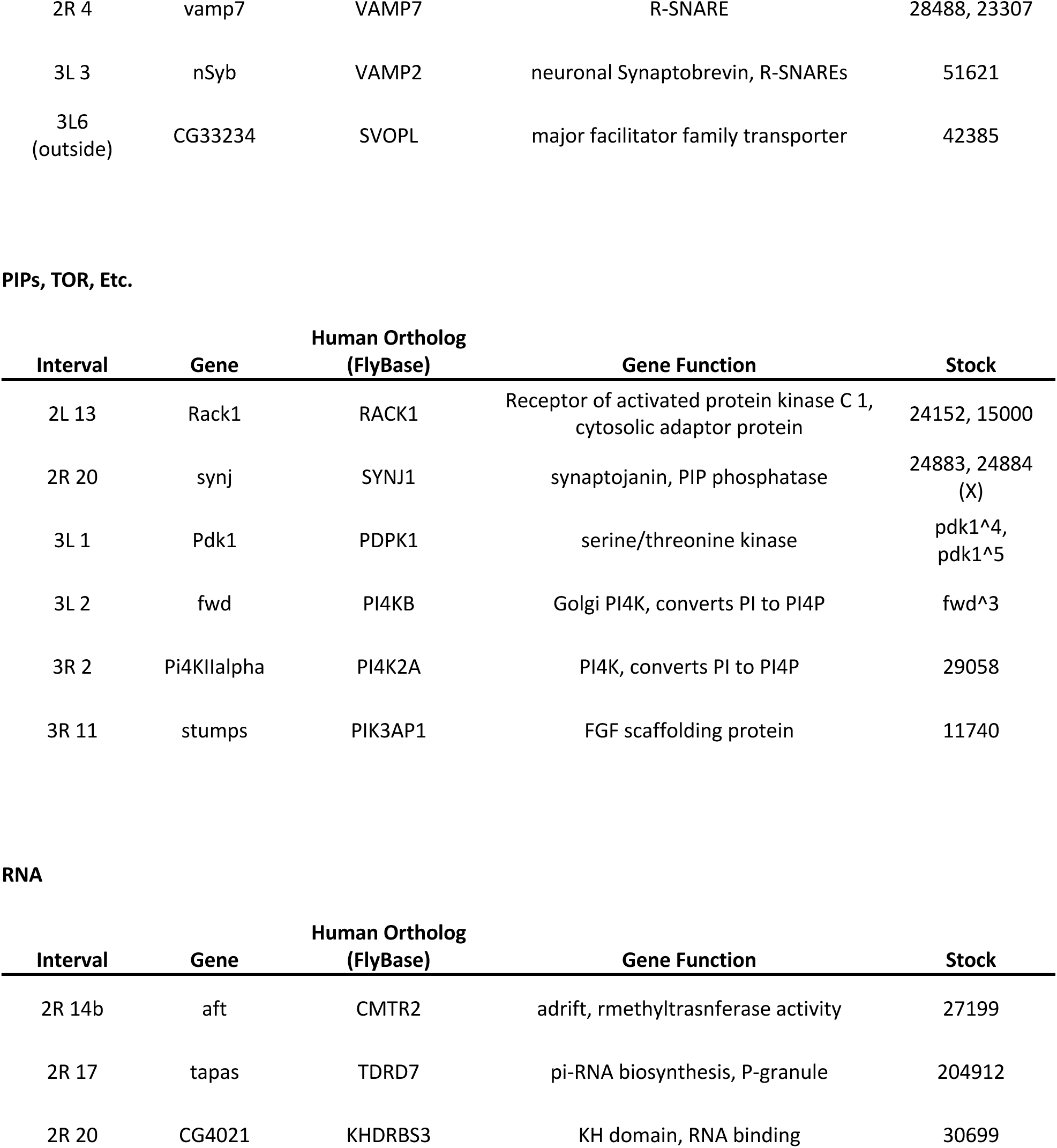
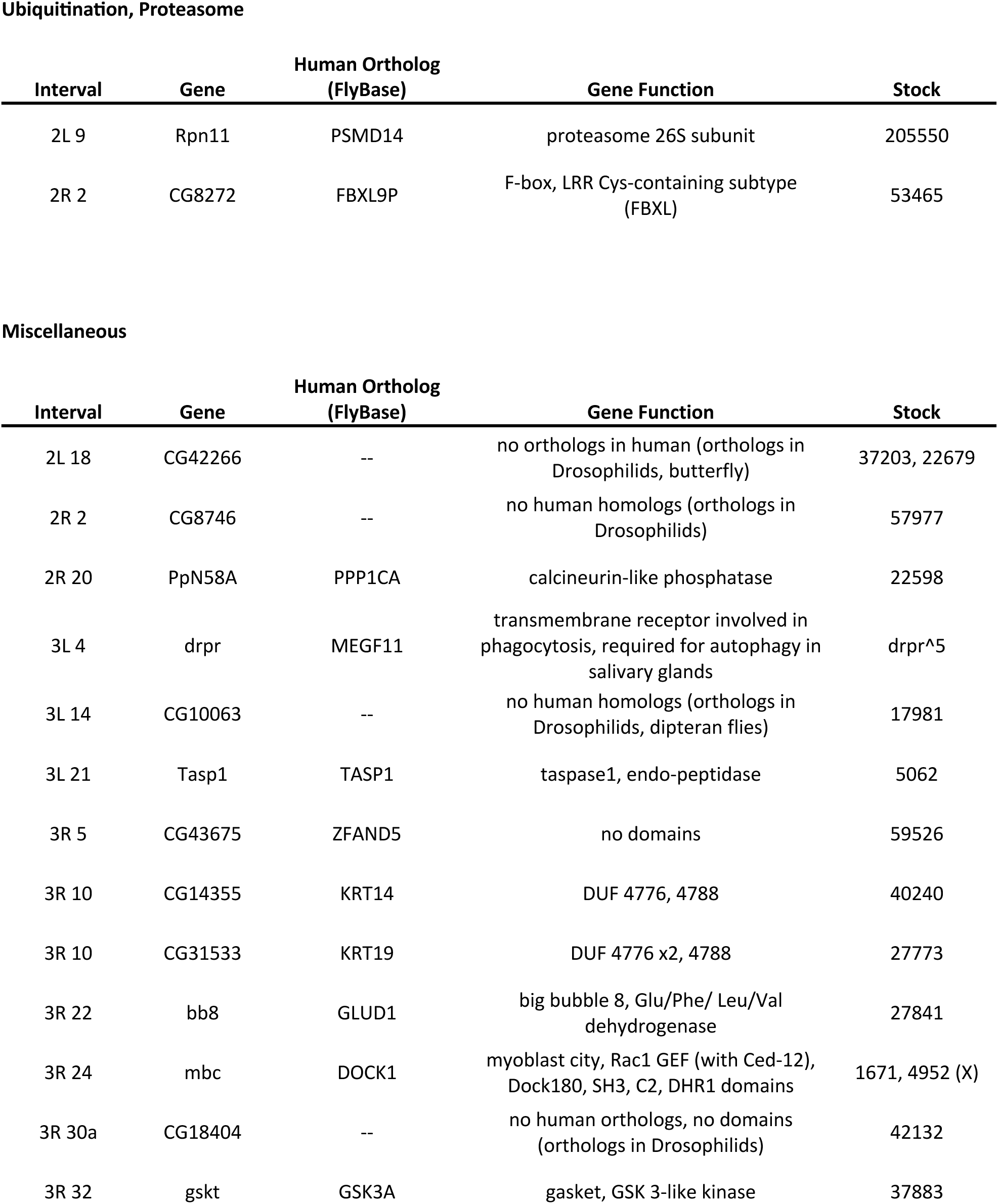

